# Shared landscape, divergent genomes: Comparative population genomics of co-occurring flora in an isolated Australian mountain range

**DOI:** 10.64898/2026.11.09.765275

**Authors:** Simón E. Lobos, Collin W. Ahrens, Paul D. Rymer, Kathryn A. Hodgins, Adam D. Miller

**Affiliations:** School of Life and Environmental Sciences, Deakin University, Warrnambool, VIC 3280, Australia; Department of Biology, School of Science, RMIT University, Melbourne, Victoria, Australia; Cesar Australia, 95 Albert St, Brunswick, VIC 3056, Australia; Hawkesbury Institute for the Environment, Western Sydney University, Richmond, NSW 2753, Australia; School of Biological Sciences, Monash University, 25 Rainforest Walk, Clayton, VIC 3800, Australia

**Keywords:** Comparative population genomics, reduced representation genome sequencing, geographic isolation, environmental isolation, landscape conservation

## Abstract

Understanding how landscape heterogeneity shapes genomic variation is key to conserving biodiversity and evolutionary potential. Yet conservation genomics is typically conducted at the single species level, limiting our ability to identify shared genomic patterns or understand how co-occurring taxa differ in their response to common environmental gradients. We assessed spatial patterns of genomic variation and their associations with geographic and environmental features among populations of nine co-existing plant species with contrasting life-histories and ecological traits across Gariwerd, a threatened biodiversity hotspot in south-eastern Australia. All species exhibited weak overall genetic structure, although the magnitude and spatial distribution of differentiation varied among species. Despite this variation, broadly consistent spatial trends were apparent, with higher genomic divergence largely concentrated in central zones and higher genomic diversity localised in southern regions of the landscape. Significant associations between genetic differentiation and geographic distance were observed in three species, while another species showed associations with environmental distance. These findings demonstrate that coexisting species can show overlapping spatial patterns in genomic variation while differing markedly in their associations with geographic and environmental variation across the same landscape. This comparative approach provides a basis for spatial prioritisation and conservation planning, while highlighting the complex eco-evolutionary processes operating across heterogeneous landscapes.

## 1. Introduction

Landscape features and environmental gradients play a central role in shaping the ecological and evolutionary processes that structure biodiversity. Spatial variation in abiotic conditions, including geography and climate, as well as biotic interactions and habitat characteristics, can influence species distributions and patterns of genetic variation within and among populations (Lee & Mitchell-Olds, 2011; Tscharntke et al., 2012; Fine, 2015). These patterns emerge through the interaction of key microevolutionary processes, including gene flow, genetic drift and natural selection, which operate across environmental and spatial gradients to shape genomic variation (Wang & Bradburd, 2014; Balkenhol et al., 2017)However, the relative influence of these processes can differ considerably within and among species, depending on their life-history traits, dispersal capacity, genomic architecture and ecological characteristics (Albecker et al., 2021; Chen et al., 2023). Understanding how these processes interact across heterogeneous landscapes is particularly important for conservation biology, as changes in habitat connectivity and environmental conditions can alter the evolutionary trajectories and the adaptive capacity of populations under ongoing environmental change.

Population genomics approaches provide a powerful framework for investigating the evolutionary processes that shape genetic variation between natural populations and for informing biodiversity conservation (Luikart et al., 2003; Breed et al., 2019). Specifically, genomic data allow us to assess the neutral evolutionary forces driving population structure, particularly the interplay between gene flow and genetic drift (Gaitán-Espitia & Hobday, 2021). Physical landscape features commonly act as barriers to dispersal, restricting gene flow and accelerating divergence among populations via genetic drift (Slatkin, 1987). Prolonged periods of reduced functional connectivity can reduce genomic diversity, increase inbreeding risk and lower fitness in small, isolated populations (Münger et al., 2026). Conversely, high levels of gene flow can maintain genomic diversity and facilitate adaptation by introducing new genomic variants, although extensive gene flow can disrupt local adaptations and enhance risks of maladaptation (Hernández-García et al., 2015). Genomic differentiation may also result from adaptive divergence, where selection favours genetic variants suited to local conditions, leading to differences in allele frequencies and phenotypes across environmental gradients (Kawecki & Ebert, 2004). Together, these processes shape spatial patterns of genetic diversity and differentiation, reflecting the complex eco-evolutionary dynamics operating across heterogeneous landscapes. Characterising these patterns and their underlying environmental drivers can identify zones of elevated diversity, pronounced divergence, or reduced connectivity, directly supporting spatial prioritisation and management (Franks et al., 2014).

In plants, pollen movement and seed dispersal are the primary mechanisms driving gene flow within and among populations (Browne et al., 2018). These dispersal mechanisms maintain functional connectivity across landscapes, helping to preserve genome-wide diversity and long-term evolutionary potential (Krauss et al., 2009). However, physical terrain and climatic gradients can either facilitate or constrain connectivity, shaping spatial patterns of genetic differentiation among populations (Manel et al., 2003; Balkenhol et al., 2017). Landscape genomic approaches provide a powerful framework for disentangling these complex environmental influences by combining genomic and environmental data with spatial modelling to quantify how landscape and climate features shape contemporary patterns of genetic variation (Balkenhol et al., 2017). Within this analytical framework, neutral markers provide an important foundation by assessing genome-wide correlations with geographic distance (Wu et al., 2015; De Vriendt et al., 2017; Mastretta-Yanes et al., 2018; Jordan et al., 2023) and environmental gradients (Jiang et al., 2019; Omondi et al., 2024), ultimately identifying areas of evolutionary significance (Endo et al., 2015; Guan et al., 2019; Yu et al., 2019).

Regions with high species richness often harbour unique evolutionary lineages that have arisen from prolonged reproductive isolation, driven by physical barriers to gene flow or divergent selection across local environmental gradients (Widmer & Lexer, 2001). A prime example is Gariwerd-Grampians National Park (hereafter “Gariwerd”) in central-western Victoria, Australia. Covering approximately 1,672 km², Gariwerd is an isolated island of native vegetation supporting approximately 1000 plant species, almost a third of the state’s indigenous flora (Enright et al., 1994). The region is geographically diverse, characterised by steep environmental gradients relating to elevation, soil composition and aridity (Arrowsmith & Inbakaran, 2002). Gariwerd is also recognised as a centre of endemism and putative refugium during past climate oscillations, likely serving as an important recruitment source for recolonising adjacent regions (Pollock et al., 2013). However, prolonged drought and high temperatures have driven several major bushfires that have severely impacted large areas of the landscape over the last 150 years (Beilin & Reid, 2015). These trends are expected to become exacerbated as the region experiences hotter, drier conditions in the coming decades (Wilkie, 2020). Although the region’s flora and fauna are well described, it remains unexplored how the fine-scale landscape features interact with divergent life histories to structure genetic variation, population connectivity, and local adaptation.

In this study, we assessed spatial patterns of genetic divergence and diversity, as well as to what extent genetic structure correlates with landscape and environmental features among co-occurring plant species with diverse life-histories and ecological features. Specifically, we examined nine unrelated plant species including trees, shrubs and a grass-tree using genome-wide SNP datasets generated through reduced genome representation sequencing. We tested whether a common landscape produces consistent spatial patterns of genomic variation or whether species exhibit distinct genomic associations with geographic and environmental features. This comparative approach distinguishes shared spatial patterns of genomic variation among coexisting species from species-specific associations with landscape heterogeneity. In doing so, it provides a framework to guide spatial prioritisation and conservation efforts aimed at preserving endemism and evolutionary potential in climate sensitive landscapes

## 2. Materials and Methods

### 2.1 Species selection and sample design

Plant species composition surveys were conducted at 147 locations from August to September 2021 (late winter/early spring; Figure S1). We selected nine widespread, sympatric plant species to provide a comparative framework spanning contrasting growth forms, life-histories and ecological characteristics (Table S1). These consisted of trees (*Banksia marginata* Cav., *Callitris rhomboidea* R.Br. ex Rich., *Eucalyptus obliqua* L’Hér.), shrubs (*Correa reflexa* (Labill.) Vent., *Epacris impressa* Labill., *Grevillea aquifolium* Lindl., *Leptospermum continentale* Joy Thomps., *Stenanthera conostephioides* Sond.), and a grass-tree species (*Xanthorrhoea australis* R.Br.; Table S1). For simplicity, species will be hereafter referred to by their genus only. In addition, this study focuses on non-model organisms with diploid genomes, none of which have an available reference genome (Rice et al., 2015).

Samples were collected from 30 sites that represented most of the range of the nine species across Gariwerd (Figure 1; Table S2). Additionally, we designated “subpopulations” with population names followed by letters to make the population of origin easy to identify. At each site, we collected leaf tissue from 10–15 plants per population per species from October 2021 to December 2022. We set a minimum sampling distance of 10m between individuals to minimise the chance of sampling siblings or close relatives. Individual samples were placed into coffee filter papers and then immediately stored into silica gel for rapid drying. Geographic coordinates were recorded at each site using a handheld Garmin GPSMAP 78 GPS device (Garmin Ltd, Schaffhausen, Switzerland). Leaf tissue was stored at room temperature and in the dark. A dry weight subsample of leaves (∼ 10–15 mg) was prepared for DNA extraction from six individuals per population per species (1602 individuals in total; range 158–188; SD 10.4; Table S2).

**Figure 1.**
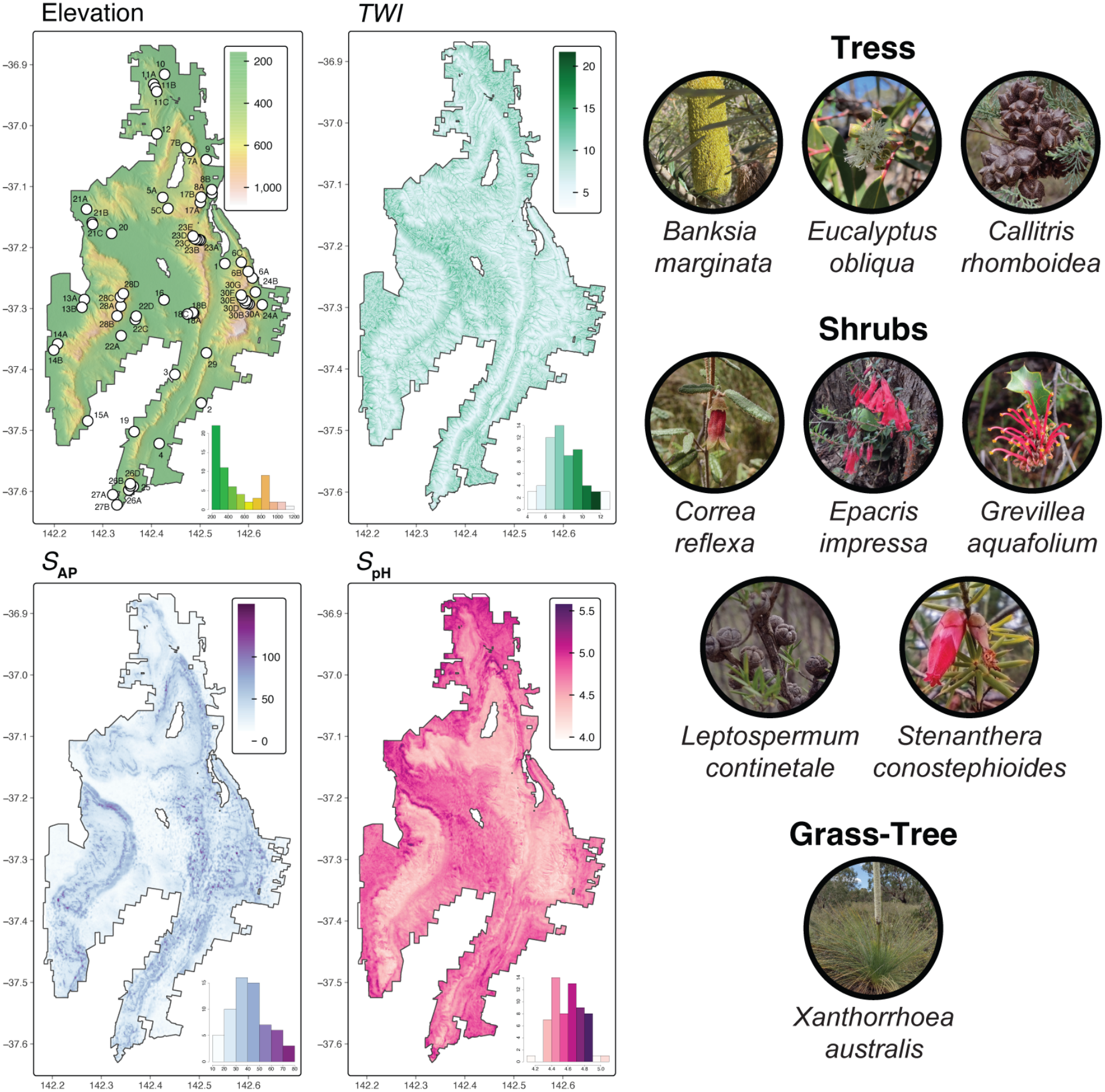
Maps showing the spatial variation of four environmental variables across Gariwerd: elevation (m), *TWI*–topographic wetness index, *S*_AP_–available phosphorus (mg/kg), and *S*_pH_–soil pH in water and CaCl_2_. Small white circles on the elevation map indicate sampling locations (see Table S2 for site details). Inset bar plots in the lower corner of each map illustrate the frequency distributions of extracted values across Gariwerd for each environmental variable. The right panel illustrates the nine plant species.

### 2.2 Total Genomic DNA Extraction and Reduced Genome Representation Sequencing

Silica-dried leaf samples were sent to the Diversity Arrays Technology sequencing facility (DArT Pty Ltd, Canberra, Australia, www.diversityarrays.com) for DNA extraction, library preparation, sequencing and SNP genotyping using proprietary DArTseq^TM^ technology. DArTseq^TM^ is a reduced genome representation sequencing method that enables genome-wide SNP genotyping without requiring a reference genome (Kilian et al., 2012) and has been widely applied across non-model plant species (e.g., *Eucalyptus*: Steane et al., 2015; Jordan et al., 2017; Rutherford et al., 2018; *Banksia*: Bradbury et al., 2019; Robins et al., 2021, 2023; Wilson et al., 2022; *Leptospermum*: Binks & Byrne, 2022). Genomic representations were generated using complexity-reduction restriction enzyme combinations optimised for each taxon: *PstI* and *HpaII* were used for *Banksia* and *Eucalyptus*, whereas *PstI* and *MseI* were applied to all other species. Following restriction digestion, specialised adaptors were ligated to the digested DNA, and ligated fragments were amplified via PCR in a thermal cycler. After PCR, amplified products were standardised, pooled and sequenced on an Illumina HiSeq 2500 platform. Libraries were sequenced at two marker densities: low-density (DArTseqLD) for *Correa*, *Leptospermum* and *Stenanthera*, and high-density (DArTseqHD) for the remaining six species (the higher-density assay was selected where greater marker density was required for a planned follow-up study). Sequences were processed using the proprietary DArT analytical pipeline (DArTsoft14) to remove low-quality reads, identify SNP markers, and assign final genotypes to each sample. To assess technical repeatability across genotype calls and library preparation, 30% of samples were resequenced as technical replicates. Additionally, technical replicates were used to assess scoring reproducibility and remove markers with low reproducibility, while testing for the Mendelian distribution of alleles was used to discriminate true allelic variants from paralogous regions and remove sequencing errors (Sansaloni et al., 2011; Kilian et al., 2012).

### 2.3 SNP quality control filtering

Further filtering of SNP loci was undertaken separately for each species using the R package *dartR* v.2.7.2 (Mijangos et al., 2022) in R v.4.2.2 (R Core Team, 2024; <https://www.rproject.org/>). Genotype data were imported as genlight objects using the R package *adegenet* v.2.1.10 (Jombart, 2008). Before SNP filtering, we screened for genetically related individuals because including clones or close relatives can bias population genetic estimates (O’Donnell et al., 2023). Pairwise genetic similarity was calculated from the proportion of shared alleles per pair of individuals using the function ‘gl.propShared’ in *dartR*. Genetic thresholds for clone assignment were calibrated using pairwise shared allele frequencies among samples for which technical replicates were available. Individuals showing 95–99% genetic similarity were initially assigned to clone groups. These assignments were subsequently evaluated using a neighbour-joining tree generated in the R package *ape* v.5.8 (Paradis & Schliep, 2019), which was also used to detect misidentified species or cross-species contamination. We retained only one representative from each clone group for downstream filtering. After relatedness screening, we removed loci with a call rate of <80%, and individuals with >20% missing genotypes across loci because substantial missing data can distort downstream analyses (Marandel et al., 2020). The remaining SNPs were filtered using a minor allele frequency (MAF) threshold of 0.05. Additionally, a Hamming distance threshold of 0.20 was applied to remove highly similar and potentially redundant loci.

### 2.4 Population genomics

For each species, we estimated population-level genomic differentiation and within-population genomic diversity. We retained at least two individuals per population, as small sample sizes can yield reliable estimates of genomic differentiation when based on a sufficiently large number of biallelic SNPs (≥1500) (Willing et al., 2012; Nazareno et al., 2017; Zbinden et al., 2023). Genomic differentiation was quantified using the Weir & Cockerham (1984) estimator of *F*_ST_, with global and pairwise estimates calculated using the ‘wc’ and ‘pairwise.WCfst’ functions, respectively, in the R package *hierfstat* v.0.5-11 (Goudet, 2005). Uncertainty was assessed using 95% confidence intervals (CIs) generated from 100 bootstrap replicates across loci using the same package, with global and pairwise CIs calculated using the ‘boot.vc’ and ‘boot.ppfst’ functions, respectively. *F*_ST_ values were considered statistically significant when their 95% CIs did not overlap zero. Genomic diversity was characterised using mean within-population expected heterozygosity (*H*_E_), observed heterozygosity (*H*_O_), and Wright’s inbreeding coefficient (*F*_IS_), calculated using the ‘basic.stats’ function in *hierfstat*. Distributions of *F*_ST_ and the three genetic diversity metrics were visualised using violin and scatterplots generated with *ggplot2* v.3.4.4 (Wickham, 2011) and pairwise population *F*_ST_ matrices were visualised as heatmaps using the R package *pheatmap* v.1.0.12 (Kolde, 2019).

Spatial patterns of population genetic structure were estimated using discriminant analysis of principal components (DAPC) in the R package *adegenet* v.2.1.10 (Jombart, 2008). DAPC is a model-free multivariate method that summarises genetic differentiation between clusters into uncorrelated variables using principal components analysis (PCA). Subsequently, a discriminant analysis (DA) is performed on the retained principal components (PCs) to maximise the between-group genetic variation while minimising within-group variation. DAPC does not make assumptions regarding the evolutionary model and Hardy-Weinberg equilibrium or linkage disequilibrium (Jombart et al., 2010). First, we calculated the highest percentage cumulative variance and lowest Bayesian Information Criterion (BIC) score to determine the most suitable clustering solution (*k* value) with the ‘find.clusters’ function. The optimal number of clusters was indicated by an elbow curve of BIC values as a function of *k* (Figures S2–S10). Next, we retained the optimal number of principal components by computing the α-score with the ‘optim.a.score’ function, as too many may result in unstable membership probabilities (Jonker et al., 2013). After determining the optimal number of PCs was determined, a second DAPC analysis was conducted using this value. As none of the analyses by species retrieved *k* > 1, further analyses were not performed.

### 2.5 Mapping patterns of genetic divergence and diversity

Genomic landscapes were generated to examine spatial patterns of genomic divergence and diversity and their correspondence with landscape features across Gariwerd. Single-species genomic surfaces were generated using the Genetic Landscapes GIS Toolbox in ArcGIS 10.8.1 (ESRI, Redlands, CA, USA; Vandergast et al., 2011). These surfaces were based on pairwise population genetic divergence (*F*_ST_) and the proportion of polymorphic loci. The proportion of polymorphic loci was calculated with the function ‘cal_prop_poly’ in the *snpR* v.1.2.8 R package (Hemstrom & Jones, 2023). To facilitate comparisons among species, we replaced the geographic coordinates of “subpopulations” with population coordinates of origin. The module creates a network of sampling locations connected to their nearest neighbours with non-overlapping edges. The midpoints between each connected edge were mapped, and genetic divergence and diversity values were assigned to these points. Single-species surfaces for each metric were then interpolated from the midpoints into a continuous surface (i.e., “genetic landscapes”) using an Inverse Distance Weighted (IDW) algorithm (power = 2, variable search radius with 12 points, ∼10m grid cell size), which assumes that the influence of the mapped variable decreases with distance from the sampled location (Vandergast et al., 2011). Spatial concordance among species was subsequently assessed using the “Multiple Species Genetic Landscape” tool, with individual surfaces standardised to a 0 to 1 scale to ensure equal weighting across species. Finally, multi-species genomic divergence and diversity rasters were summarised as average and variance surfaces.

### 2.7 Environmental data collection

Environmental data for each sampling location were retrieved from open-source GIS databases in raster format to represent bioclimatic, edaphic, and hydrological gradients (Figure 1). The environmental characteristics of each sampling location were quantified using a set of six relevant predictors, based on past work defining them as eco-physiologically important (Williams et al., 2012; Guerin et al., 2022; Mokany, McCarthy, et al., 2022; Westerband et al., 2023). Four bioclimatic variables, including annual mean temperature (*T*_MA_), maximum temperature of the warmest month (*T*_MAX_), annual precipitation (*P*_MA_) and precipitation of the driest month (*P*_MIN_) were obtained from the WorldClim v.2.06 database (Fick & Hijmans, 2017) as gridded files at 30 arc-second resolution (∼ 1 km^2^) for the years 1970–2000. Two soil variables, available phosphorus–*S*_AP_ and soil pH in water and CaCl_2_–*S*_pH_, were obtained from the TERN Soil and Landscape Grid of Australia database (https://data.csiro.au/; Grundy et al., 2015) at 3 arc-second resolution (∼ 90 x 90 m pixels) and 0–5cm soil depth, as plant roots are mainly distributed in surface soils and because there are very strong correlations across depths (Luo et al., 2021). Finally, to account for the landscape’s hydrological characteristics, we used the topographic wetness index (*TWI*), as a proxy for soil moisture conditions. *TWI* represents local topographic influence on hydrological processes based on catchment area and local slope inclination (Kopecký et al., 2021). *TWI* derived at 10m resolution from digital elevation models (DEMs) was downloaded through the CSIRO Data Access Portal (https://data.csiro.au/collections; Gallant & Austin, 2015). To account for the differing spatial resolutions, all environmental rasters were calibrated and resampled to a uniform spatial resolution of 1 arc-second ( ∼30 × 30m grid cell size) using the *raster* v.3.6-26 (Hijmans, 2023) and *rgdal* v.1.6-7 packages in R (Bivand et al., 2006-2023).

Collinearity among all predictor variables was estimated by applying a hybrid approach that involves quantifying the variance inflation factor (VIF) (Marquardt, 1970) and pairwise correlation tests implemented in the ‘vifcor’ function in the R package *usdm* v.2.1-7 (Naimi et al., 2014). This approach uses a stepwise procedure to quantify VIF for all variables, calculate correlations between variable pairs, and identify the pair with the highest correlation value. A correlation coefficient threshold of 0.6 was used, following the recommendations of Sales et al. (2021). Strong collinearity was detected among bioclimatic predictor variables, as well as between bioclimatic predictors and *S*_pH_ (Figure S11). Consequently, all bioclimatic variables were excluded from subsequent analyses due to collinearity issues and their coarse spatial resolution (∼1km), which fails to capture local microclimate and microhabitat information.

### 2.8 Genetic isolation by distance and environment

We assessed the relative importance of geographic distance (isolation-by-distance; IBD) and environmental dissimilarity (isolation-by-environment; IBE) in explaining the genomic differentiation for each species using Generalised Dissimilarity Modelling (GDM; Ferrier et al., 2007). GDM is based on community-level modelling algorithms that use matrix regression to estimate nonlinear relationships between pairwise genetic dissimilarity, geographic distance, and environmental variables (Ferrier et al., 2007). We selected GDM over traditional distance-based methods like the Mantel test, which has been criticised for high false-positive rates with non-independent data in landscape genomics analyses (Legendre et al., 2015). Briefly, GDM fits a linear combination of monotonic I-spline functions to each predictor to quantify (1) the variation in the rate (the shape of each spline) of genetic turnover at different localities along each environmental and geographic gradient and (2) the curvilinear relationships between genetic distance and environmental and geographic distances (Fitzpatrick & Keller, 2015). The highest point reached by each spline indicates the magnitude of total genetic change along that gradient, reflecting the relative importance of that predictor in influencing genetic turnover while keeping all other variables constant.

Prior to model fitting, the variables were converted into site-pair format using the ‘formatsitepair’ function In the R package *gdm* v.1.5.0-9.1 (Fitzpatrick et al., 2022). Pairwise *F*_ST_ matrices were used as the biological response variable, and predictor data sets were assembled with geographic coordinates of each sampling location and the three environmental variables described above. To examine how turnover rate and intensity vary along predictor gradients across the nine species, we used the ‘isplineExtract’ function to extract I-spline values for species associated with at least one predictor. We then evaluated model statistical significance and predictor importance using the ‘gdm.varimp’ function (1000 permutations within the same package. The function compares the original model’s deviance explained against models built from randomly permuted predictor data across sites. The p-value represents the proportion of permuted runs that equalled or outperformed the observed data. Variables above the significance threshold (typically *p* ≥ 0.05) can then be pruned from the model (Mokany, Ware, et al., 2022). The *p*-value represents the proportion of permuted runs that equalled or outperformed the observed data. Variables above the significance threshold (typically *p* ≥ 0.05) can then be pruned from the model (Mokany, Ware, et al., 2022).

## 3. Results

### 3.1 SNP quality control filtering

A total of 708,421 biallelic SNPs (range 18,960–190,206; SD 63,366.6) were initially identified from 1577 individuals (range 151–188; SD 12.8) across nine species. Of these, 70,129 SNPs (range 18,960–31,698; SD 7211.3) were generated using DArTseqLD and 638,292 SNPs (range 20,212–190,026; SD 60,396.6) using DArTseqHD (Table S3). Following filtering for call rate, missingness, minor allele frequency and paralogous loci, 1299 individuals (range 106–174; SD 21.2) were retained with a final dataset of 90,330 (range 757–28,654; SD 9,839.8) high quality SNP loci for subsequent populations genomic analyses (DArTseqLD: range 757–4326; SD 1805.5 SNPs and DArTseqHD: range 1938–28,654; SD 10,044.3 SNPs). These final datasets represented 20–30 sampling locations per species (mean = 26.8; SD 3.3; Table S3), providing broad spatial coverage across Gariwerd.

### 3.2 Population genomics

Measures of overall genomic differentiation revealed moderate structure (*F*_ST_ = 0.0527–0.0891) in four of nine species (e.g., *Stenanthera*, *Correa*, *Banksia* and *Grevillea*), while weak levels (*F*_ST_ = 0.0336–0.0474) were found in the remaining five (e.g., *Callitris*, *Leptospermum*, *Epacris*, *Xanthorrhoea* and *Eucalyptus*; Table S4 and Figure 2). Across collection localities, pairwise comparisons pairwise *F*_ST_ comparisons displayed a notable incidence of high divergence (*F*_ST_ > 0.1) within *Epacris* (0.3%; Figure 2 and Figure S5), *Correa* (0.5%; Figure 2 and Figure S4), *Banksia* (7%; Figure 2 and Figure S2) and *Grevillea* (8%; Figure 2 and Figure S7). Notably, most collection localities within these species (except *Epacris*) showed moderate differentiation, accounting for ∼77% of all pairwise comparisons (Figure 2). Within *Epacris*, pairwise *F*_ST_ between collection localities was generally low (≥75%), with the greatest differentiation between locations at the southern (Mt Abrupt–26A) and northern (Pohlners Rd–11C; FST = 0.1646) ends of Gariwerd (Figure 2 and Figure S5). In contrast, low pairwise differentiation predominated across *Leptospermum* (65%; Figure 2 and Figure S8), *Callitris* (66%; Figure 2 and Figure S3), *Eucalyptus* (88%; Figure 2 and Figure S6) and *Xanthorrhoea* (90%; Figure 2 and Figure S10), indicating a low level of population differentiation among locations, consistent with genomic connectivity and weak regional structuring across the landscape. Finally, collection localities within *Stenanthera* were moderately differentiated (52%; Figure 2 and Figure S9).

**Figure 2.**
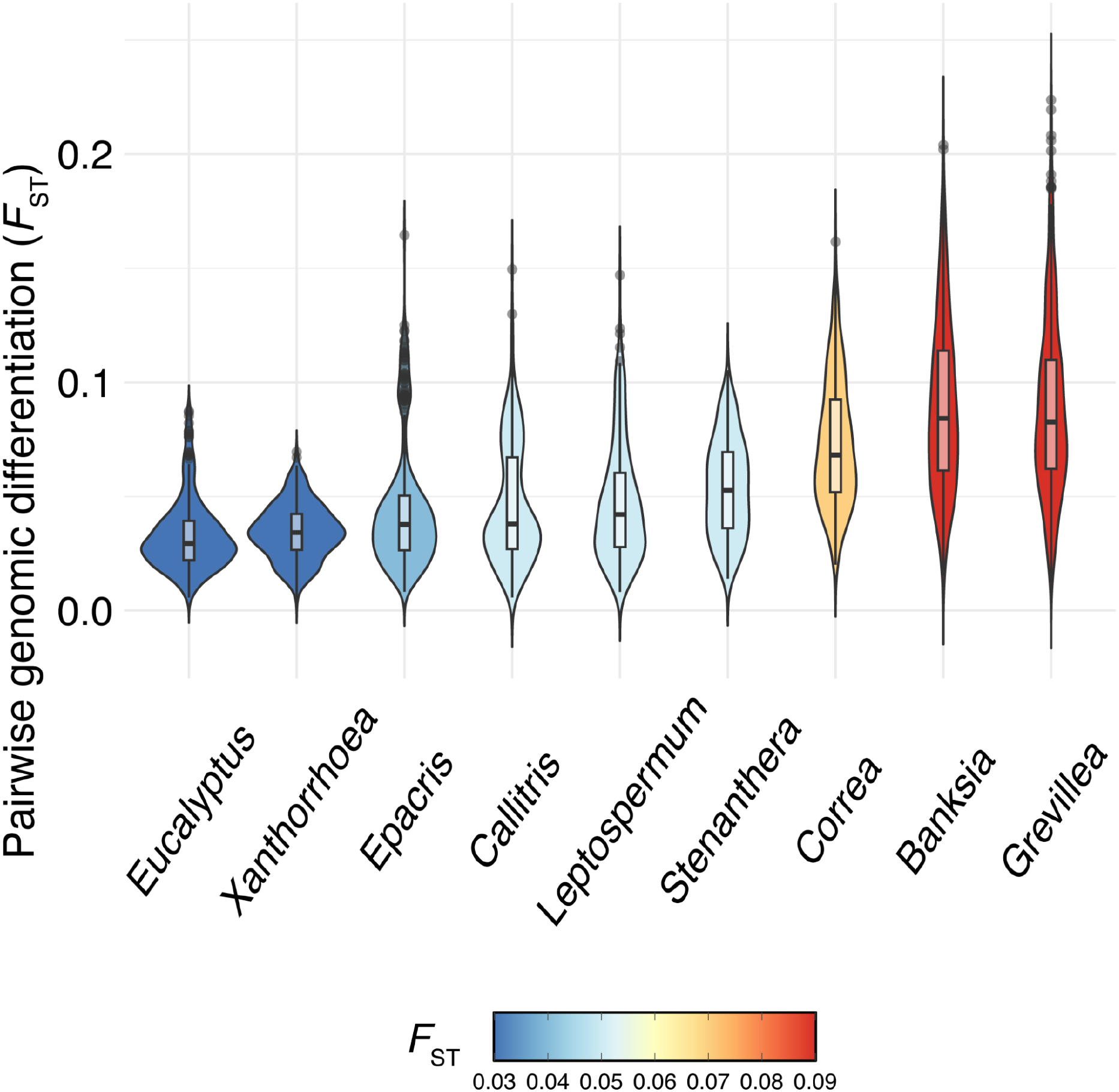
Violin plots showing the distribution of pairwise genomic differentiation (*F*_ST_) within each of the nine species. For the boxplot, the lower horizontal bound shows the first quartile (25%) and the upper horizontal bound shows the third quartile (75%), with the black horizontal line as the median. Outliers are represented as small black circles. The colour gradient represents the overall genomic differentiation value (*F*_ST_).

At the species level, observed heterozygosity was relatively low (∼15%) in *Callitris*, *Correa*, *Epacris*, and *Grevillea*. In contrast, *Leptospermum*, *Banksia*, *Eucalyptus*, *Stenanthera*, and *Xanthorrhoea* showed moderate levels, averaging approximately 19% observed heterozygosity (Table S4). Overall, observed heterozygosity was consistently lower (*H*_O_ = 0.0716–0.2407) than expected heterozygosity (*H*_E_ = 0.1445–0.3118) across collection localities, except for five populations of *Correa* (e.g., Henham Track–3, Grampians Rd–4, Mt Zero Rd–10, Pohlners Rd–12 and Harrops Track–13A; Figure 3 and Figure S4). Measures of heterozygosity within *Leptospermum*, *Stenanthera* and *Eucalyptus* exhibited little variation among collection localities, whereas *Grevillea*, *Xanthorrhoea* and *Epacris*, showed greater inter-population variation. Finally, *Callitris* and *Correa* showed the lowest overall genetic diversity estimates (Figure 3; Table S4).

**Figure 3.**
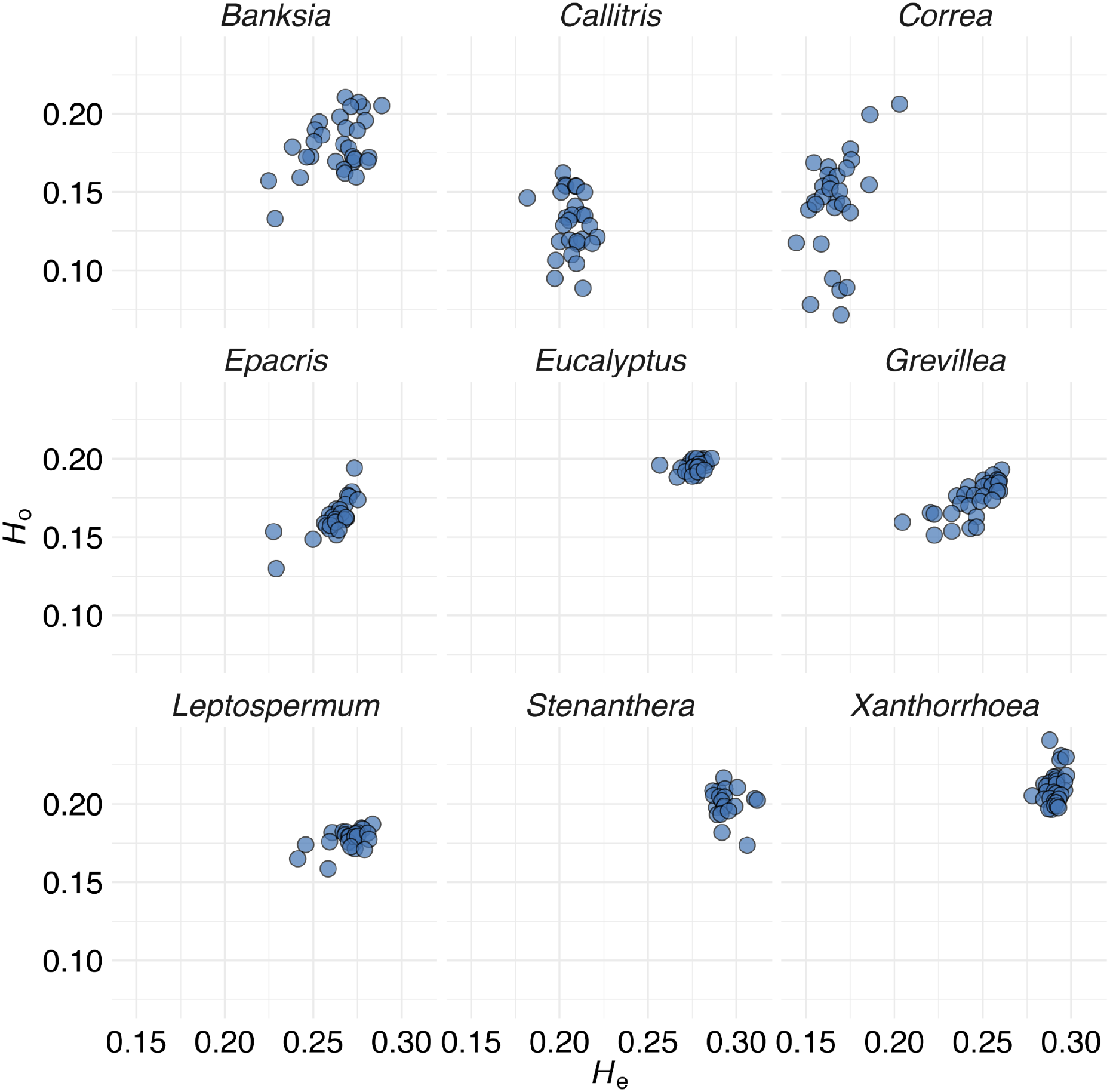
Comparison of within-population genetic diversity for each of the nine species. Observed heterozygosity (*H*_O_; y-axis) is plotted as a function of expected heterozygosity (*H*_E_; x-axis) for each collection locality (represented by circles).

DAPC analysis found no indication of discrete genetic clusters across the nine plant species in Gariwerd (Figures S2–S10). The Bayesian Information Criterion (BIC) values increased consistently with increasing k, indicating that k = 1 was the best-supported solution.

### 3.3 Mapping patterns of genetic divergence and diversity

The averaged genomic divergence landscapes for all taxa ranged from 0.17 to 0.56 ( mean = 0.37), indicating clear latitudinal genomic differences across the northern, central, and southern reaches of Gariwerd (Figure 4A). Concurrent low divergence patterns were identified primarily on the east side of the northern zone (182 km^2^; 11% of the analysed area) and the northern half of the southern zone (412 km^2^; 26%) (Figure 4A). This second region of low divergence extends from the western boundary of the southern Victoria Range, down to the western side of the Victoria Valley, spreading across the Serra Range (e.g., Mirranatwa Gap, Mt Lang, Teddy Bear Gap) and up to the base of the Mt William Range (Figure 4A). Conversely, high divergence occurred in the centre of Gariwerd, extending from Mt Thackery through Victoria Valley to the summit of Mt Lubra (144 Km^2^; 9%; Figure 4A). A second region of high divergence encompasses two small areas located in the south-eastern extreme of the Victoria Range (Bullawin Rd–15) across the Victoria Valley at the level of Bryan Swamp (∼17 Km^2^; 1%; Figure 4A). High-divergence locations across Gariwerd are genetically distinct from all other locations and from one another. Variance among divergence landscapes was typically low across most parts of the study area, highlighting broad interspecific congruence. However, variance was greater at a few locations in our sampling distribution because each species showed unique high-divergence patterns (Figure 4B).

**Figure 4.**
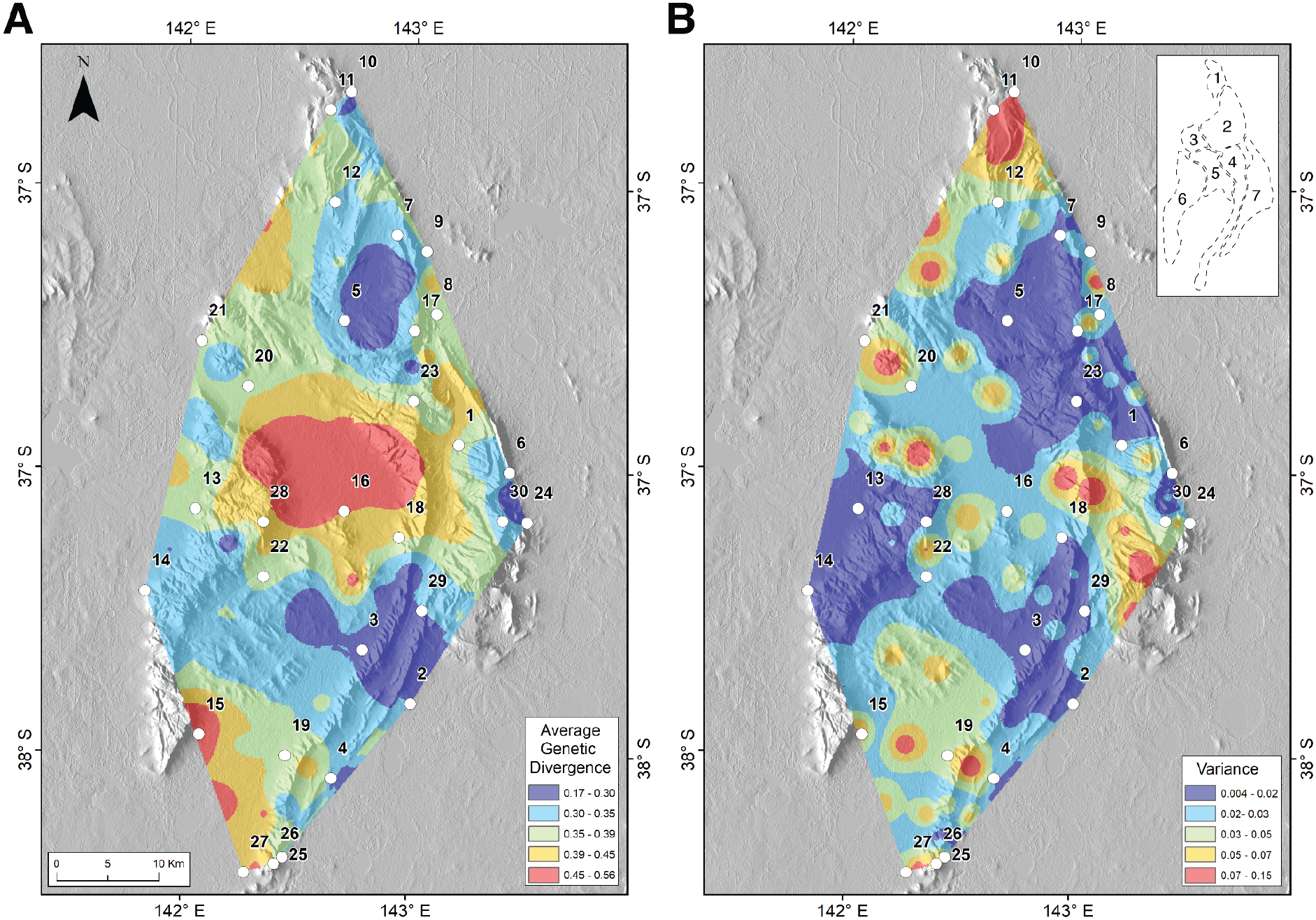
Multiple species genetic divergence landscapes: (A) average surface from nine plant species across Gariwerd and (B) variance surface among species landscapes. Values are classified into five groups, with the highest values in red and the lowest values in blue. All other colours indicate the intermediate level of divergence. White circles represent sampling locations (see Table S2 for location names). The top-right inset map outlines the regions of Gariwerd (1: Mount Stapylton Area; 2: Mount Difficult Range; 3: Mackenzie River Area; 4: Serra Range; 5: Victoria Valley; 6: Victoria Range; and 7: Mount William Range).

Across species, the surfaces of average genetic diversity ranged from 0.47 to 0.82, with a median of 0.66. The highest diversity occurred in the southern zone of Gariwerd, mostly spread across the eastern side of this region and covering 22% (354 km^2^) of the study area (Figure 5A). Notable peaks of high diversity were recorded at the populations Redman Rd–6, Mt Zero Rd–10, Pohlners Rd–11, Lodge Rd–20 and Glenelg River Rd–22 (Figure 5A). Conversely, genetic diversity was relatively low across the centre of the Mt Difficult Range and the eastern side of the Mackenzie River area, towards the valley between Mt Rosea and Mt William (257 km^2^; 16%; Figure 5A). Similarly, low genetic diversity was also recorded at the southernmost extreme of Gariwerd between populations Grampians Rd–25, Mt Abrupt–26 and Victoria Valley Rd–27 (55 km^2^; 3%; Figure 5A). Variation among diversity landscapes was generally low across Gariwerd (0.002–0.17; *<u>x</u>* = 0.07); however, the highest variance occurred around two sampling locations in the northeast region (Mackenzie Rd–5 and Mt Rosea–23; Figure 5B).

**Figure 5.**
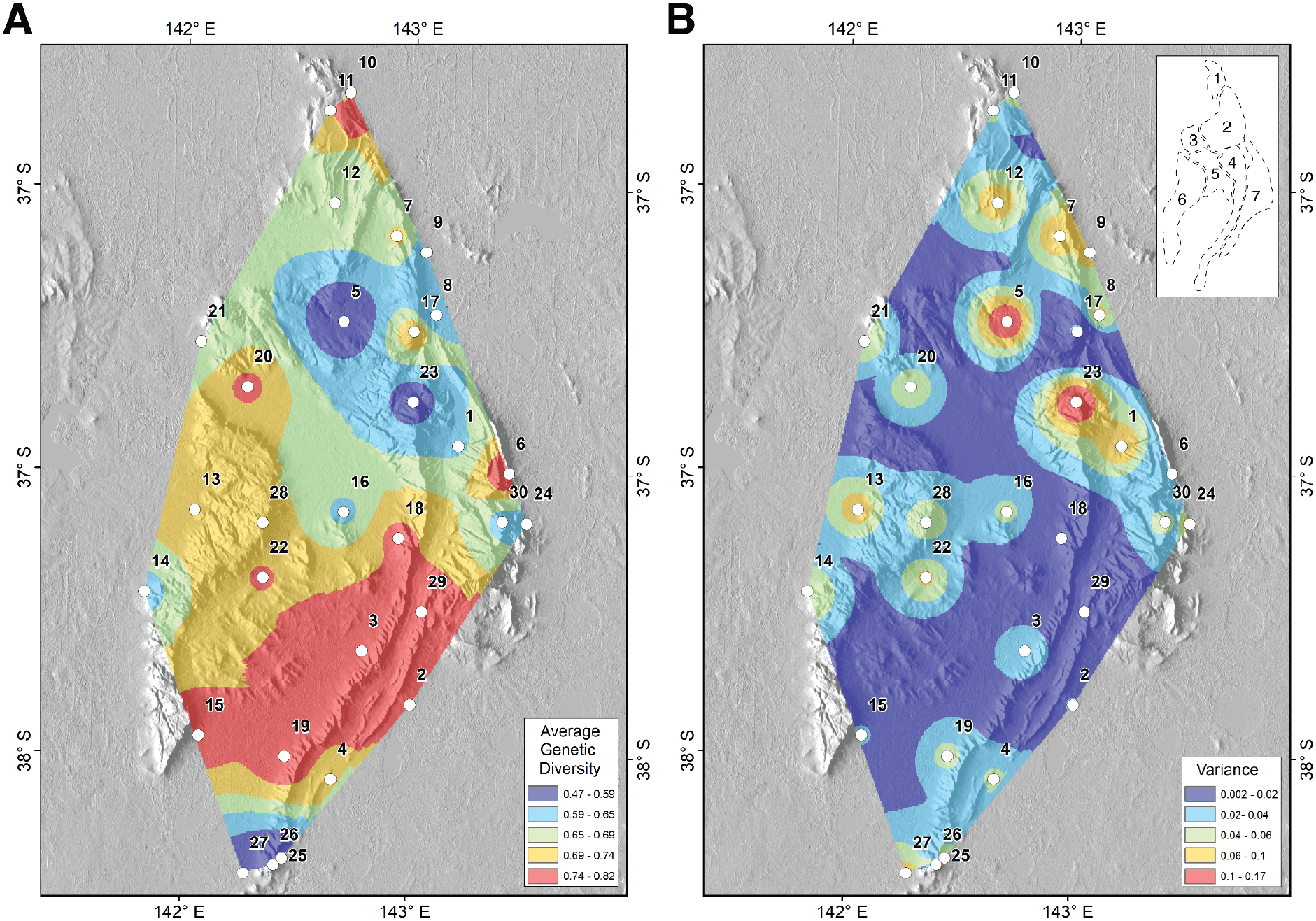
Multiple species genetic diversity landscapes, measured using the proportion of polymorphic loci: (A) average surface from nine plant species across Gariwerd and (B) variance surface among species landscapes. Values are classified into five groups, with the highest values in red and the lowest values in blue. All other colours indicate intermediate diversity levels. White circles represent sampling locations (see Table S2 for location names). The top-right inset map outlines the regions of Gariwerd (1: Mount Stapylton Area; 2: Mount Difficult Range; 3: Mackenzie River Area; 4: Serra Range; 5: Victoria Valley; 6: Victoria Range; and 7: Mount William Range).

### 3.5 Genetic isolation by distance and environment

The analysis of the effects of geographic distance and environmental dissimilarity on genetic differentiation revealed statistically significant patterns only for *Correa* (*p* = 0.005) and *Stenanthera* (*p* < 0.001), explaining approximately 23.79% and 28.11% of overall deviance, respectively, (Figure 6). Although the overall model for *Grevillea* was not statistically significant, geographic distance did influence genomic turnover (*p* = 0.005). This indicates that the relationship between genomic turnover and the variables analysed was not uniform across the studied taxa. For these two significant models, geographic distance was significantly correlated with pairwise *F*_ST_, accounting for 7.78% (*p* = 0.037) for *Correa* and 92% (*p* < 0.001) for *Stenanthera* of relative predictor importance, respectively (Figure 6A). Looking at the fitted patterns across taxa, *Grevillea* and *Stenanthera* showed the steepest genomic turnover over short geographic distances (between ∼2 and ∼20 km), although *Grevillea* displayed a more moderate pattern across the gradient (Figure 6A). Conversely, for *Correa*, genomic differentiation increased with geographic distance, but more gradually (Figure 6A). Environmental distance was significantly associated with genomic differentiation only in *Stenanthera*, with *S*_pH_ accounting for 4.91% (*p* < 0.001) of the deviance explained. For *Stenanthera,* genetic differentiation increased logarithmically with *S*_pH_, accelerating before reaching a plateau around pH ∼4.7 (Figure 6D).

**Figure 6.**
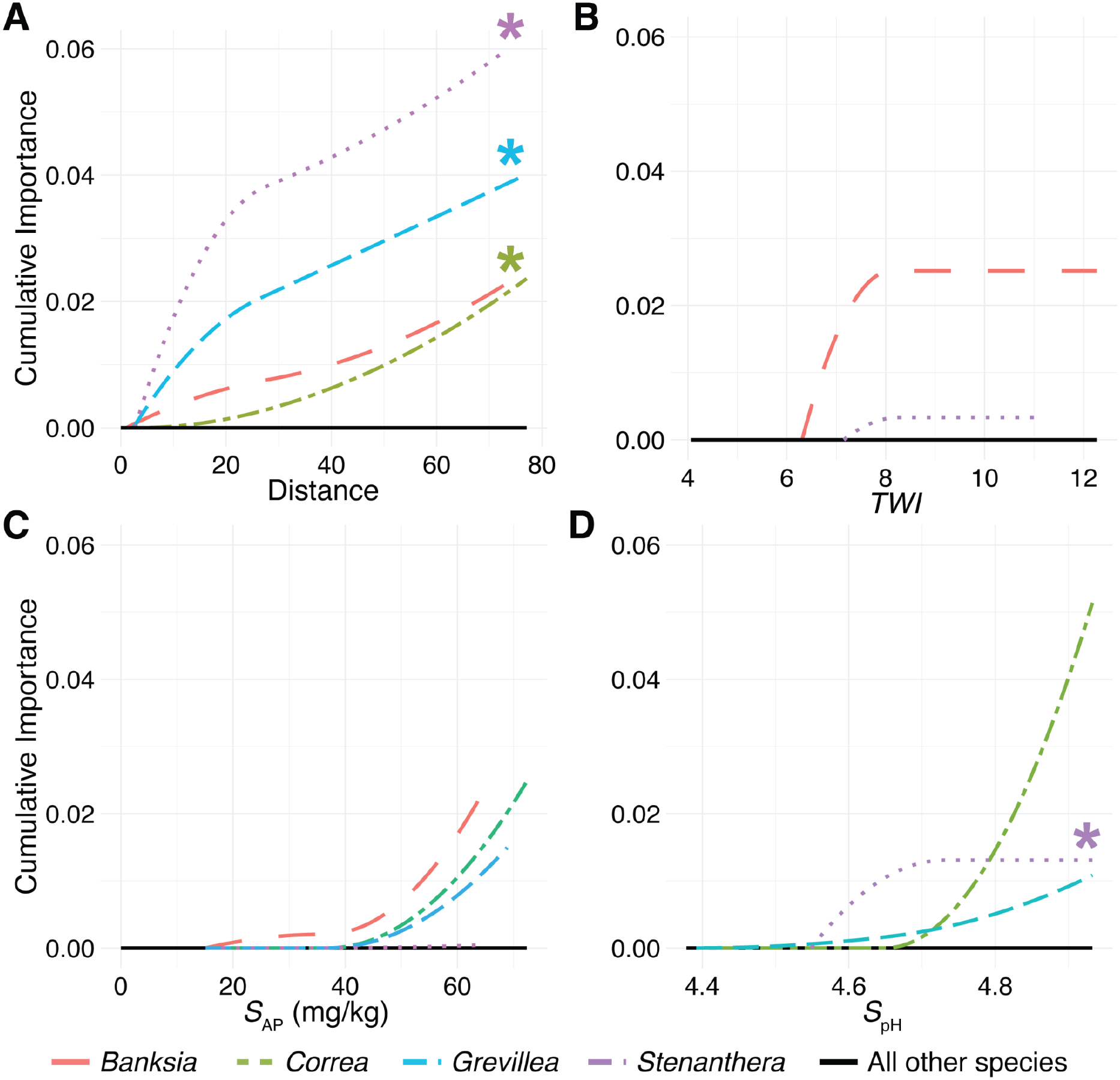
Generalised dissimilarity model-fitted I-splines showing genetic turnover across (A) distance and (B–D) environmental distance for *Banksia marginata*, *Correa reflexa*, *Grevillea aquifolium* and *Stenanthera conostephioides*. Significant relationships are denoted by the (*) symbol. The maximum height of each curve indicates the total amount of genetic turnover along the gradient when holding all other variables constant. The I-spline’s shape informs about the rate of genetic turnover. Variable abbreviations are *TWI*–topographic wetness index, *S*_AP_–available phosphorus, and *S*_pH_–soil pH in water and CaCl_2_.

## 4. Discussion

Evaluating the impact of spatial environmental heterogeneity on genomic variation across co-occurring species is important for isolating broad landscape-scale processes from species-specific responses and for guiding conservation management. Using a comparative approach across nine sympatric plant species, we found both shared and individual-taxa patterns of genomic variation. Contrary to our expectation that topographic and microclimatic heterogeneity would drive strong population structure, all species exhibited relatively low overall genomic structure. However, genomic differentiation and diversity showed broadly concordant spatial patterns across species, with relatively high differentiation in central areas and relatively high genomic diversity in southern areas of Gariwerd. Associations between genomic differentiation and geographic or environmental distance varied widely among species, indicating that a common landscape context can produce broadly concordant spatial patterns of genomic variation without imposing a common mechanism of differentiation across species. Together, these findings suggest that coexisting species share fine-scale spatial trends across this portion of their range landscape despite differing in their IBD and IBE patterns, however similarities in habitat requirements, dispersal, or other ecological aspects may also contribute.

### 4.1 Spatial concordance in genomic differentiation

The high spatial correspondence in genomic differentiation across species with contrasting ecology and life histories suggests that landscape elements and environmental factors strongly influence the spatial distribution of genome-wide variation. The highest mean divergence across species occurred at the south-western limit of Gariwerd, extending longitudinally through farmland and the Bryan Swamp wetland complex between the Victoria and Serra ranges (Figure 4A). Here, open grassland and habitat fragmentation likely constrain functional connectivity (Lozada-Gobilard et al., 2021). A second area of elevated differentiation occurred in central Gariwerd, within the Victoria Valley and the flanking slopes and valleys between the three parallel sandstone ranges, where variation in elevation and habitat may similarly constrain movement. In contrast, lower differentiation occurred across parts of the northern and southern zones where fewer obvious ecological or topographic barriers may facilitate connectivity. This finding suggests that landscape composition and configuration can impose a shared spatial template on genomic differentiation.

Compared to previous studies within Gariwerd, our genetic estimates were generally lower than those reported in both single-species (Evans et al., 2001) and multi-species studies (Pollock et al., 2013). Such differences likely reflect differences in molecular marker systems, species-specific evolutionary histories and sampling regimes used to capture extant diversity (Mariette et al., 2002; Landguth et al., 2012). Moreover, given that these species are naturally widespread across Australia with several isolated populations, fine-scale selection pressures or historic isolation in diverse environments may drive localised divergence (Evans et al., 2001; Ng et al., 2014; Richardson et al., 2014; Senior et al., 2022).

### 4.2 Spatial concordance in genomic diversity

A complementary pattern emerged for genomic diversity, which was consistently concentrated in south-eastern Gariwerd, across the eastern portion of the Victoria Range and Mount William region. A similar pattern was previously observed in *Eucalyptus baxteri*, where intensive fine-scale sampling identified high chloroplast haplotype diversity associated with sheltered east-facing slopes and valleys in the southern Serra Range (Pollock et al., 2013). The recurrence of high diversity across this region, supported by broader landscape genetic studies (Vandergast et al., 2013; Wood et al., 2013; Guan et al., 2019), suggests that microclimatically stable or sheltered areas act as long-term refugia that promote the maintenance and persistence of genomic variation. Such areas may function as important reservoirs of evolutionary diversity and, potentially, as sources of genetic variation for surrounding populations through ongoing dispersal and recruitment. Consequently, landscape-scale genomic analyses can help identify potential evolutionary refugia of high conservation importance.

### 4.3 Species-specific drivers of genomic differentiation

The broad spatial concordance among species contrasts with the limited evidence that geographic or environmental distance independently explains genomic differentiation within individual taxa. Geographic distance was associated with genomic differentiation in *Correa, Grevillea and Stenanthera*, whereas evidence for IBE was detected only in *Stenanthera*, driven by soil pH. The limited evidence for IBE across the remaining species may reflect differences in ecological associations among taxa, unmeasured abiotic drivers of differentiation, or insufficient regional environmental variation. In a heterogeneous landscape like Gariwerd, unmeasured microclimate, substrate, or moisture variation may influence connectivity and selection at finer spatial scales than our data reflect. Ultimately, these results demonstrate that similar spatial patterns of genomic differentiation can arise across the landscape despite differences in underlying ecology, genomic architecture and evolutionary processes.

Dispersal-related traits provide one potential explanation for these species-specific patterns because they directly influence gene flow. For example, *Grevillea* produces relatively heavy, gravity-dispersed seeds, which may restrict seed-mediated gene flow and contribute to the observed IBD pattern. Conversely, pollen-mediated dispersal may be particularly important in bird-pollinated species. *Stenanthera* and *Correa* are predominantly pollinated by the territorial New Holland Honeyeater (*Phylidonyris novaehollandiae*) (Paton & Ford, 1977), whose foraging ecology and aggressive territorial behaviour may restrict pollen movement over larger distances (McFarland, 1996). These species-specific differences in dispersal pathways provide a plausible ecological basis for variation in IBD strength among taxa. The mechanisms underlying IBE may be similarly complex, as environmental gradients can simultaneously drive spatially varying selection (Ellegren & Sheldon, 2008), shape habitat suitability (Ortego et al., 2012), flowering phenology (Dahlgren et al., 2007), and alter pollinator dynamics (Papanikolaou et al., 2017; Glück et al., 2022). Such effects could alter patterns of pollen-mediated gene flow across the landscape, strengthening differentiation where environmental transitions coincide with reduced pollinator movement or divergent selection (Schiestl & Johnson, 2013; Wessinger, 2021). In *Stenanthera*, soil pH was strongly associated with genomic differentiation, which may be ecologically relevant given that soil pH shapes the structure and development of Ericaceae mycorrhizal associations (Lambers et al., 2021). Previous work identified differences in root-associated fungi between *Stenanthera* populations in Gariwerd and populations elsewhere in Victoria (McLean & Lawrie, 1996), providing a potential mechanism linking local soil conditions to population differentiation. Overall, differences in dispersal capacity, reproductive biology, pollinator interactions, and local environmental responses may explain why IBD and IBE relationships vary among taxa. Rather than operating through a single overarching mechanism, the landscape imposes a common spatial template of genomic variation. At the same time, individual species follow divergent micro-evolutionary trajectories driven by species-specific traits.

### 4.4 Implications for conservation and management

Characterising spatial patterns of genomic variation across multiple species provides a practical framework for spatially explicit conservation planning in heterogeneous landscapes like Gariwerd. Although Gariwerd is recognised as a biodiversity hotspot and long-term refuge for Australian flora and fauna (Pollock, Bayly, et al., 2015; Pollock, Rosauer, et al., 2015), genomic information has not been widely integrated into its management. Our identification of broadly concordant patterns of genomic differentiation and diversity across multiple species offers a basis for prioritising areas for protection, restoration and connectivity. Small, isolated populations persisting in environmentally distinct habitats may be particularly vulnerable to environmental change because of reduced genetic diversity, adaptive capacity and recruitment potential (Willi et al., 2006; Mosblech et al., 2011). Areas of high genomic differentiation highlight where landscape-scale environmental features impede functional connectivity, signalling priority regions for targeted restoration. Conversely, areas of high diversity represent vital reservoirs of evolutionary variation that warrant long-term protection and can aid recovery after major disturbances, such as fire, by supplying seed, pollen or recolonising individuals. Ultimately, this comparative approach establishes landscape-scale priorities that extend beyond the requirements of individual taxa.

Quantifying gene flow along environmental and elevational gradients underpins climate-adaptive management. High-elevation populations face rapid environmental shifts, where functional connectivity from lower-elevation populations may influence their capacity to persist (Sexton et al., 2014). Our multispecies framework provides a baseline to identify populations needing urgent evaluation, particularly where elevated differentiation or low diversity coincides with geographic isolation. Distinguishing locally adapted populations from those shaped primarily by drift or restricted gene flow requires combining complementary analyses of adaptive genomic divergence with quantitative experiments. For populations that are both locally adapted and isolated under climatic shifts, assisted gene flow could be considered to enhance adaptive potential and resilience (Aitken & Whitlock, 2013; Sacristán-Bajo et al., 2025). Together, these findings extend conservation planning beyond individual taxa by identifying landscape-scale genomic relationships that support the retention, functional connectivity, and adaptive potential of entire plant communities

## Author Contributions

A.D.M. received funding. A.D.M. and S.E.L. conceived and designed the study, with input and feedback from C.W.A. and P.D.R. A.D.M. and S.E.L. conducted the field work. S.E.L. performed the bioinformatic and statistical analyses. S.E.L., A.D.M. and K.A.H. drafted the manuscript. All authors read, edited and approved the final manuscript.

## Acknowledgements

We thank Erlania, Emily Armstrong, Zach Clark, Harry Coleman, Jessica Fish, Owen Holland, and Daniel Vairo from Deakin University for their support during sample collections. A special thanks to Bill Weatherly (Friends of Forgotten Woodlands Inc.), Adam Merrick (Trust for Nature) and Simon Heyes (La Trobe University) for assistance with species and site selection.

## Ethical Statement

Permission to collect tissue samples was provided by DEECA under permit number 10010169 and adhered to all local guidelines.

## Benefit Sharing Statement

This research contributes to the conservation of flora in Gariwerd by openly sharing its findings with DEECA to inform and support local management efforts. Additionally, all data and results are publicly accessible, as detailed in the Data Availability Statement.

## Data Availability Statement

All data, along with the code required to reproduce the reported results are openly accessible via the Dryad Data Repository and the GitHub repository (https://github.com/selobos/Conservation-Genomics). Raw sequence reads have been deposited in the National Centre for Biotechnology Information (NCBI) Sequence Read Archive (https://www.ncbi.nlm.nih.gov/sra/PRJNA1484297).

