## Supplemental Material for "Shared landscape, divergent genomes: Comparative population genomics of co-occurring flora in an isolated Australian mountain range"

**Table of Contents:**

| **Table S1.** Summary of life history traits for the species examined in the study. | Page 2−3 |
| --- | --- |
| **Table S2.** Total number of individuals per species within each sampling location (n=30) used for the analyses. | Page 4−8 |
| **Table S3.** Total number of SNPs retained for population genomic analyses. | Page 9 |
| **Table S4.** The results of the *F*-statistics analysis and genetic diversity for each species. | Page 10 |
| **Figure S1.** Map of the spatial distribution of the 147 survey locations across Gariwerd. | Page 11 |
| **Figure S2−S10.** (A) Modelling of the genetic differences among “populations” using a DAPC.  (B) Comparison of genetic diversity parameters. (C) Heatmap of pairwise *F*_ST_ between all  sampling location populations (Weir & Cockerham *F*_ST_). | Page 12−20 |
| **Figure S11.** Heatmap of Pearson correlation coefficients showing relationships between  extracted environmental predictors across sampling locations for generalised  dissimilarity modelling (GDM). | Page 21 |
| **Figure S12.** Genetic divergence landscape of the nine species from Gariwerd. | Page 22 |
| **Figure S13.** Genetic diversity landscape of the nine species from Gariwerd. | Page 23 |

**Table S1.** Summary of life history traits for the species examined in the study.

| **Species** | **Family** | **Mating**  **System** | **Seed dispersal**  **mechanism** | **Life-**  **form** | **Pollen Vector** |
| --- | --- | --- | --- | --- | --- |
| *Banksia marginata* | Proteaceae | Primarily outcrossing, but is capable of self-fertilisation^1^ | Gravity^2^ | Tree | Birds, insects and/or small non-flying mammals |
| *Callitris rhomboidea* | Cupressaceae | Predominantly outcrossing^3^ | Wind and gravity^4^ | Tree | Wind |
| *Eucalyptus obliqua* | Myrtaceae | Primarily outcrossing, but is capable of self-fertilisation^5^ | Wind-shake and gravity^6^ | Tree | Insects, bats, and/or birds |
| *Correa*  *reflexa* | Rutaceae | Primarily outcrossing, but is capable of self-fertilisation^7^ | Presumably explosive release and ants^8^ | Shrub | Bees and/or birds |
| *Epacris impressa* | Ericaceae | Primarily outcrossing, but is capable of self-fertilisation^9^ | Gravity and wind^10^ | Shrub | Birds and/or Insects |
| *Grevillea*  *aquifolium* | Proteaceae | Unknown | Presumably gravity and ants^11^ | Shrub | Birds and/or insects |
| *Leptospermum continentale* | Myrtaceae | Primarily outcrossing, but is capable of self-fertilisation^12^ | Gravity^13^ | Shrub | Insects |
| *Stenanthera conostephioides* | Ericaceae | Presumably  outcrossing^14^ | Animal-dispersed seed^15^ | Shrub | Birds |
| *Xanthorrhoea australis* | Asphodelaceae | Presumably  outcrossing^16^ | Gravity^16^ | Grass-tree | Insects, birds small non-flying mammals |

**^1^**Miller et al. (2020)

^2^Clarke et al. (2010)

^3^Liu et al. (2024)

^4^Harris & Kirkpatrick (1991)

^5^Brown et al. (1975)

^6^Facelli & Kerrigan (1996)

^7^Vivian M. Butz Huryn (1997)

^8^Auld (2001)

^9^Fripp (1982)

^10^Worth et al. (2017)

^11^Holmes et al. (2014)

^12^O’Brien (1994)

^13^Plumanns-Pouton et al. (2024)

^14^Keighery (1996)

^15^Ghazoul (2005)

^16^Lamont et al. (2004)

**Table S2.** Total number of individuals per species within each sampling location (n=30) used for the analyses. Numbers after the slash indicate the subset of individuals selected for DArTseq sequencing. Underlined numbers indicate that a specimen was sequenced twice as a technical replicate. Species abbreviations are as follows: BM, *Banksia marginata*; CR, *Callitris rhomboidea*; CX, *Correa reflexa*; EI, *Epacris impressa*; EO, *Eucalyptus obliqua*; GA, *Grevillea aquifolium*; LC, *Leptospermum continentale*; SC, *Stenanthera* *conostephioides*, and XA, *Xanthorrhoea australis*.

| **Locations** | **Code** | **Species** | | | | | | | | | **Elev.(m)** | **Lat. (ºS)** | **Lon. (ºE)** |
| --- | --- | --- | --- | --- | --- | --- | --- | --- | --- | --- | --- | --- | --- |
|  |  | **BM** | **CR** | **CX** | **EI** | **EO** | **GA** | **LC** | **SC** | **XA** |  |  |  |
| Redman Rd | 1 | 4/6 | 4/6 | 7 |  | 1/6 | 6/6 |  | 6/6 | 6/7 | 344 | -37.23 | 142.55 |
| Yarram Gap Rd | 2 | 5/7 | 5/6 | 6 | 5/7 | 5/7 | 6/7 |  | 6/7 | 6/6 | 307 | -37.46 | 142.50 |
| Henham Track | 3 | 4/6 | 6/6 | 7 | 6/6 | 6/6 | 6/6 | 6/7 | 6/6 | 6/7 | 312 | -37.41 | 142.45 |
| Grampians Rd | 4 | 4/6 | 6/6 | 6 | 4/6 | 5/6 | 6/6 | 6/6 |  | 6/6 | 259 | -37.52 | 142.42 |
| Mackenzie Rd | 5A | 3/6 |  | 6 | 3/6 | 2/6 | 6/6 | 6/6 |  | 6/6 | 454 | -37.12 | 142.42 |
|  | 5C |  | 6/6 |  |  |  |  |  |  |  | 571 | -37.14 | 142.43 |
| Redman Rd | 6A | 6/6 | 6/7 | 6 | 6/7 |  |  |  | 6/6 | 6/6 | 431 | -37.25 | 142.61 |
|  | 6B |  |  |  |  |  |  | 6/6 |  |  | 599 | -37.24 | 142.60 |
|  | 6C |  |  |  |  |  | 6/6 |  |  |  | 452 | -37.22 | 142.59 |
| Mt Difficult Rd | 7A |  | 6/6 | 6 |  |  | 6/7 | 6/7 | 3/6 | 6/6 | 534 | -37.04 | 142.48 |
|  | 7B | 6/7 |  |  |  | 6/6 |  |  |  |  | 474 | -37.04 | 142.47 |
| Mt Zero Rd | 8A | 5/6 |  |  | 6/6 | 1/6 |  | 6/6 |  | 6/6 | 248 | -37.11 | 142.52 |
|  | 8B |  | 6/6 | 6 |  |  | 6/6 |  |  |  | 249 | -37.11 | 142.52 |
| Mt Zero Rd | 9 | 6/7 | 6/6 | 6 | 6/6 | 6/6 | 6/7 | 6/6 | 3/6 | 6/6 | 248 | -37.06 | 142.51 |
| Mt Zero Rd | 10 | 6/7 | 6/7 | 6 |  | 3/6 | 6/6 |  | 6/7 | 6/6 | 228 | -36.92 | 142.43 |
| Pohlners Rd | 11A |  | 5/6 | 6 |  |  | 6/6 |  |  | 6/6 | 389 | -36.93 | 142.40 |
|  | 11B |  |  |  |  | 5/6 |  |  |  |  | 365 | -36.94 | 142.41 |
|  | 11C | 6/6 |  |  | 5/6 |  |  | 6/6 |  |  | 369 | -36.94 | 142.41 |
| Pohlners Rd | 12 | 6/6 | 5/7 | 6 | 5/6 | 6/7 | 6/6 | 6/7 | 5/6 | 6/6 | 436 | -37.01 | 142.41 |
| Harrops Track | 13A | 5/6 | 6/6 | 7 | 6/6 |  | 6/6 | 6/6 | 6/6 | 4/7 | 236 | -37.29 | 142.26 |
|  | 13B |  |  |  |  | 6/6 |  |  |  |  | 239 | -37.30 | 142.26 |
| Harrops Track | 14A | 3/6 | 6/6 | 6 | 6/7 | 5/6 |  | 6/6 | 4/6 | 6/6 | 284 | -37.36 | 142.21 |
|  | 14B |  |  |  |  |  | 6/6 |  |  |  | 266 | -37.37 | 142.20 |
| Bullawin Rd | 15A | 6/7 | 6/7 | 7 | 6/6 | 6/7 | 6/7 | 6/6 | 5/7 | 6/7 | 279 | -37.48 | 142.27 |
| Serra Rd | 16 | 5/7 | 5/6 | 7 | 6/6 | 6/7 | 6/7 | 6/6 | 6/7 | 5/7 | 264 | -37.29 | 142.43 |
| Boroka Lookout | 17A | 5/6 | 5/6 | 6 | 5/7 | 4/6 | 6/6 |  |  | 6/6 | 852 | -37.13 | 142.50 |
|  | 17B |  |  |  |  |  |  | 6/7 |  |  | 826 | -37.12 | 142.50 |
| Henham Track | 18A |  | 6/7 |  |  |  |  |  |  |  | 424 | -37.31 | 142.49 |
|  | 18B | 5/6 |  | 6 | 5/7 |  | 6/6 |  | 5/6 | 6/6 | 350 | -37.31 | 142.49 |
|  | 18C |  |  |  |  |  |  | 6/6 |  |  | 277 | -37.31 | 142.47 |
| Serra Range Fireline | 19 | 4/6 | 6/6 | 7 | 6/6 | 5/6 | 6/7 | 6/7 | 6/7 | 6/7 | 289 | -37.50 | 142.36 |
| Lodge Rd | 20 | 4/6 | 5/7 | 6 | 6/6 | 6/6 | 6/6 | 6/6 | 6/6 | 6/6 | 229 | -37.18 | 142.32 |
| Asses Ear Rd | 21A | 4/7 | 6/6 | 6 |  |  | 6/6 | 6/7 | 5/6 | 6/6 | 228 | -37.14 | 142.27 |
|  | 21B |  |  |  |  | 4/7 |  |  |  |  | 226 | -37.16 | 142.28 |
|  | 21C |  |  |  | 5/6 |  |  |  |  |  | 218 | -37.16 | 142.28 |
| Glenelg River Rd | 22A | 5/6 |  | 7 | 6/6 | 4/6 | 6/6 | 5/6 |  |  | 277 | -37.34 | 142.34 |
|  | 22C |  |  |  |  |  |  |  |  | 6/7 | 259 | -37.32 | 142.37 |
|  | 22D |  |  |  |  |  |  | 6/6 | 6/6 |  | 263 | -37.31 | 142.37 |
| Mt Rosea | 23A |  | 6/6 |  | 3/7 |  |  |  |  |  | 846 | -37.19 | 142.50 |
|  | 23B |  |  |  |  |  | 6/7 |  |  |  | 846 | -37.19 | 142.50 |
|  | 23C | 4/6 |  |  |  |  |  |  |  |  | 777 | -37.19 | 142.49 |
|  | 23D |  |  | 6 |  |  |  |  |  |  | 708 | -37.19 | 142.49 |
|  | 23E |  |  |  |  | 6/6 |  | 4/7 |  | 6/6 | 685 | -37.18 | 142.49 |
| Mitchell Rd | 24A | 5/6 | 3/6 | 7 | 5/6 | 6/6 |  | 6/6 | 5/7 | 6/7 | 333 | -37.29 | 142.63 |
|  | 24B |  |  |  |  |  | 6/6 |  |  |  | 357 | -37.27 | 142.61 |
| Grampians Rd | 25 | 3/7 | 4/6 |  | 6/6 | 5/7 | 6/6 | 5/6 |  | 5/6 | 372 | -37.59 | 142.36 |
| Mt Abrupt | 26A | 5/6 |  |  | 4/6 |  |  |  |  | 6/6 | 816 | -37.60 | 142.35 |
|  | 26B |  | 4/6 |  |  |  |  |  |  |  | 641 | -37.59 | 142.36 |
|  | 26D |  |  | 6 |  |  |  |  |  |  | 584 | -37.59 | 142.36 |
| Victoria Valley Rd | 27A | 5/6 | 5/6 | 6 |  | 6/7 | 6/6 | 4/6 | 5/6 | 6/6 | 221 | -37.61 | 142.32 |
|  | 27B |  |  |  | 6/6 |  |  |  |  |  | 270 | -37.62 | 142.33 |
| Mt Thackeray | 28A |  |  |  | 6/7 | 6/6 |  |  |  |  | 966 | -37.30 | 142.34 |
|  | 28B |  |  |  |  |  | 6/7 |  |  | 6/6 | 856 | -37.31 | 142.33 |
|  | 28C |  | 6/7 |  |  |  |  |  |  |  | 786 | -37.28 | 142.34 |
|  | 28D | 5/7 |  | 6 |  |  |  |  |  |  | 814 | -37.28 | 142.34 |
| Jimmy Creek Rd | 29 | 6/6 |  | 6 | 6/6 | 5/6 | 6/6 |  | 6/6 | 6/6 | 335 | -37.37 | 142.51 |
| Mt William | 30A | 5/6 |  |  |  |  |  |  |  |  | 1161 | -37.29 | 142.60 |
|  | 30B |  |  |  | 4/7 |  |  |  |  |  | 1076 | -37.29 | 142.60 |
|  | 30D |  |  |  |  |  |  |  |  | 6/6 | 1009 | -37.29 | 142.59 |
|  | 30E |  | 6/7 |  |  |  | 6/6 |  |  |  | 943 | -37.29 | 142.59 |
|  | 30F |  |  |  |  | 6/6 |  | 6/7 |  |  | 858 | -37.28 | 142.59 |
|  | 30G |  |  | 6 |  |  |  |  |  |  | 803 | -37.28 | 142.58 |

**Table S3** Total number of SNPs retained for population genomic analyses.

| **Species** | **Pre-filtering** | | | | **Post-filtering** | | |
| --- | --- | --- | --- | --- | --- | --- | --- |
|  | **No. SNPs** | **No. individuals** | **No. populations** | **No. clones/**  **excluded**  **ind.** | **No. SNPs** | **No. individuals** | **No. populations** |
| **High-density** |  |  |  |  |  |  |  |
| *Banksia marginata* | 20,212 | 188 | 30 | 16 | 1,938 | 145 | 30 |
| *Callitris rhomboidea* | 72,381 | 176 | 28 | 0 | 10,840 | 152 | 28 |
| *Epacris impressa* | 135,235 | 164 | 27 | 0 | 11,264 | 143 | 27 |
| *Eucalyptus obliqua* | 80,226 | 188 | 30 | 46* | 7,376 | 130 | 25 |
| *Grevillea aquifolium* | 190,026 | 182 | 29 | 0 | 23,109 | 174 | 29 |
| *Xanthorrhoea australis* | 140,212 | 188 | 30 | 4* | 28,654 | 174 | 30 |
| **Low-density** |  |  |  |  |  |  |  |
| *Correa reflexa* | 18,960 | 171 | 29 | 7 | 757 | 137 | 29 |
| *Leptospermum continentale* | 31,698 | 151 | 25 | 0 | 4,326 | 138 | 24 |
| *Stenanthera* *conostephioides* | 19,471 | 169 | 27 | 50* | 2,229 | 106 | 20 |

*****Pre-filtering individual counts include samples excluded as clonemates or misidentified species.

**Table S4** The results of the *F*-statistics analysis (overall genetic differentiation–*F*_ST_, confidence interval–CI and overall inbreeding coefficient–*F*_IS_) and genetic diversity (observed proportion of heterozygosity–*H*_O_ and expected proportion of heterozygosity–*H*_E_) for each species.

| **Species** | ***F*_ST_** | **CI** | ***F*_IS_** | ***H*_O_** | ***H*_E_** |
| --- | --- | --- | --- | --- | --- |
| *Banksia marginata* | 0.088 | 0.085–0.092 | 0.310 | 0.180 | 0.263 |
| *Callitris rhomboidea* | 0.047 | 0.046–0.049 | 0.364 | 0.130 | 0.207 |
| *Correa reflexa* | 0.070 | 0.064–0.075 | 0.139 | 0.143 | 0.167 |
| *Epacris impressa* | 0.043 | 0.041–0.044 | 0.374 | 0.163 | 0.262 |
| *Eucalyptus obliqua* | 0.034 | 0.032–0.035 | 0.290 | 0.197 | 0.278 |
| *Grevillea aquifolium* | 0.089 | 0.088–0.09 | 0.282 | 0.174 | 0.244 |
| *Leptospermum continentale* | 0.047 | 0.045–0.049 | 0.337 | 0.177 | 0.269 |
| *Stenanthera conostephioides* | 0.053 | 0.049–0.056 | 0.315 | 0.199 | 0.293 |
| *Xanthorrhoea australis* | 0.035 | 0.034–0.035 | 0.273 | 0.211 | 0.291 |

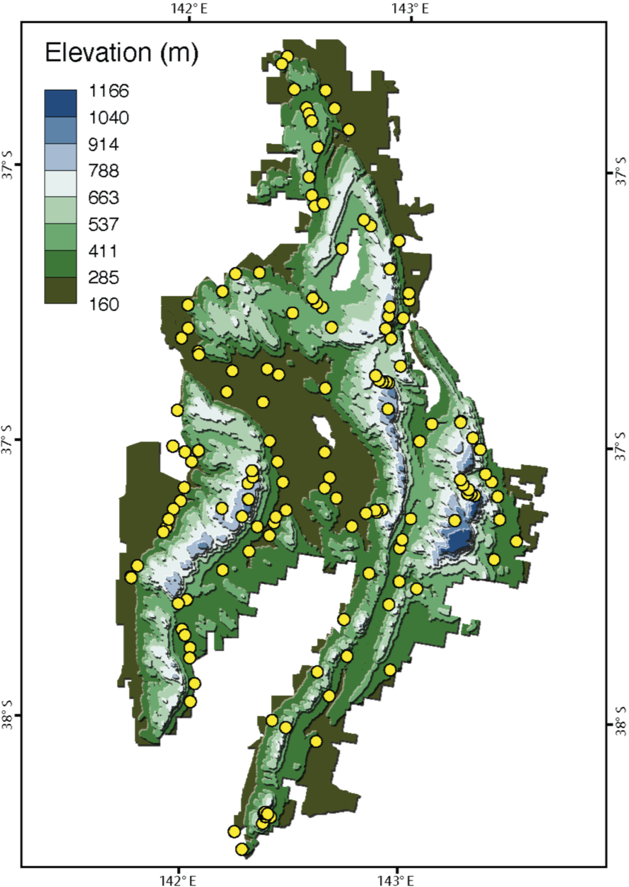

**Figure S1** Map of the spatial distribution of the 147 survey locations across Gariwerd. Contour lines represent the height of the ground above sea level (m.a.s.l.) in meters.

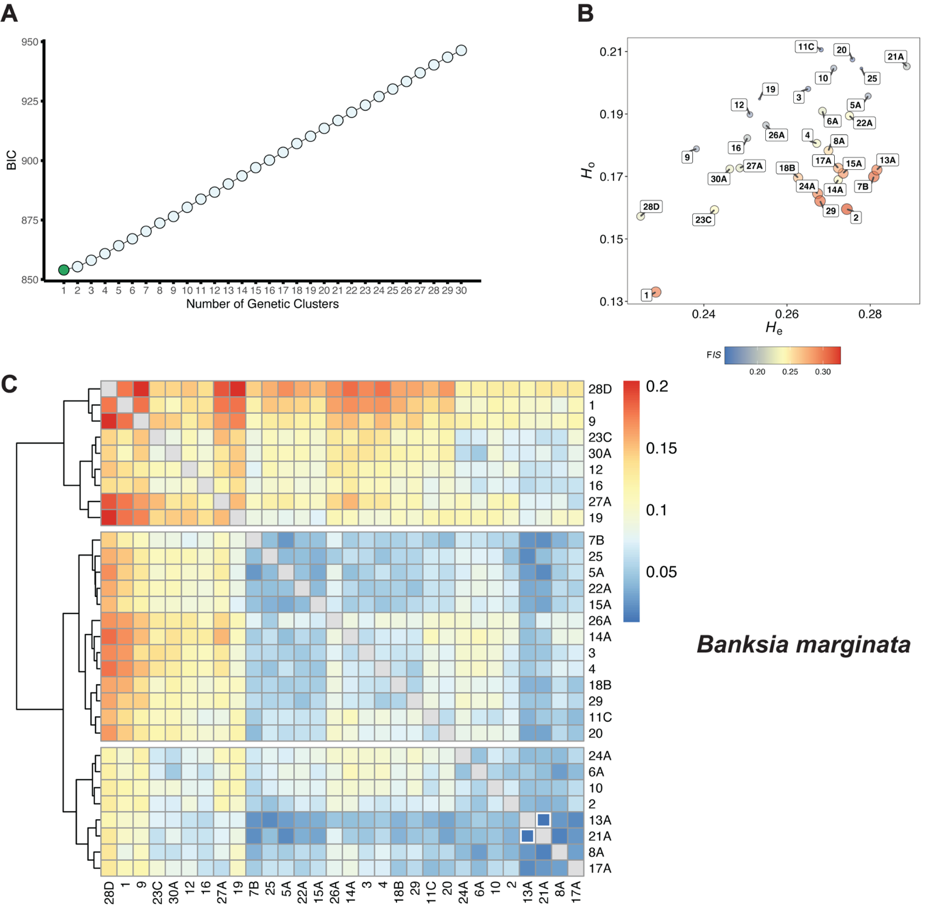

**Figure S2** (A) Modelling of the genetic differences among “populations” using a DAPC. (B) Comparison of genetic diversity parameters, where observed heterozygosity (*H*_O_; y-axis) was plotted as a function of expected heterozygosity (*H*_E_; x-axis). Locations are coloured by the overall inbreeding coefficient (*F*_IS_), and circle size is proportional to this value. (C) Heatmap of pairwise *F*_ST_ between all sampling location populations (Weir & Cockerham *F*_ST_). White frame squares reflect confidence intervals (CI) less than 0. In figures b) and c), the colour blue indicates lower estimates, and yellow and red indicate middle and higher estimates, respectively.

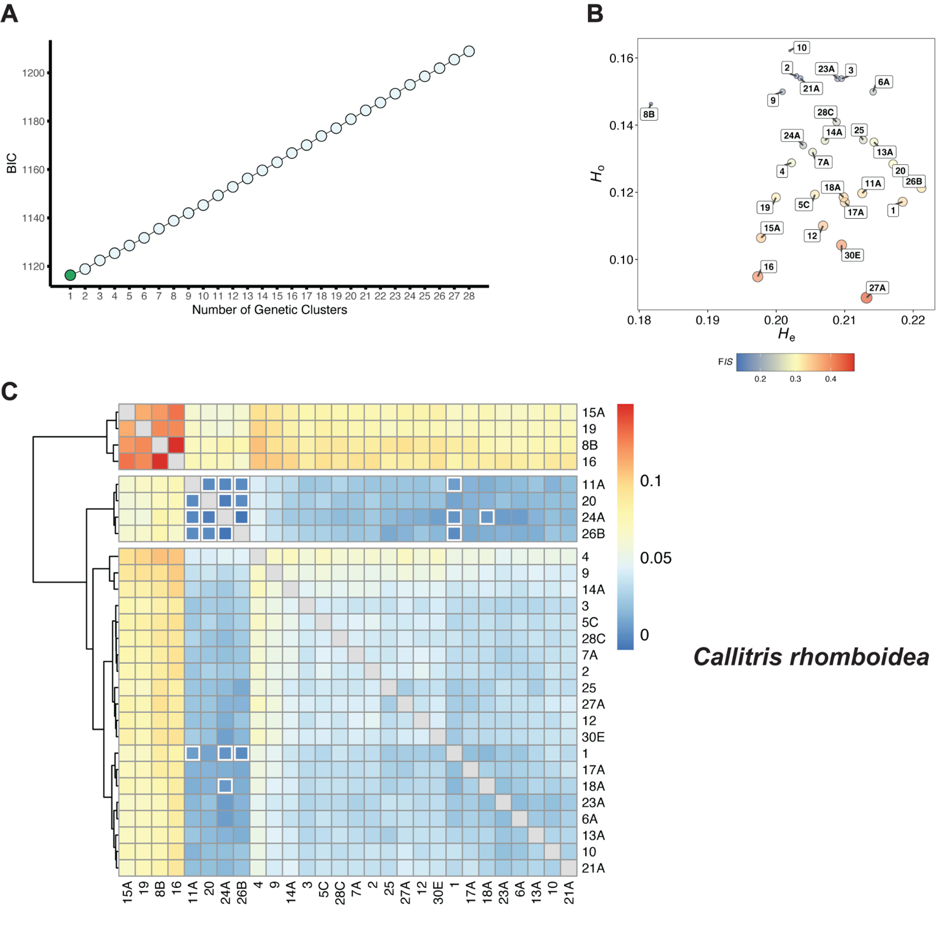

**Figure S3** (A) Modelling of the genetic differences among “populations” using a DAPC. (B) Comparison of genetic diversity parameters, where observed heterozygosity (*H*_O_; y-axis) was plotted as a function of expected heterozygosity (*H*_E_; x-axis). Locations are coloured by the overall inbreeding coefficient (*F*_IS_), and circle size is proportional to this value. (C) Heatmap of pairwise *F*_ST_ between all sampling location populations (Weir & Cockerham *F*_ST_). White frame squares reflect confidence intervals (CI) less than 0. In figures b) and c), the colour blue indicates lower estimates, and yellow and red indicate middle and higher estimates, respectively.

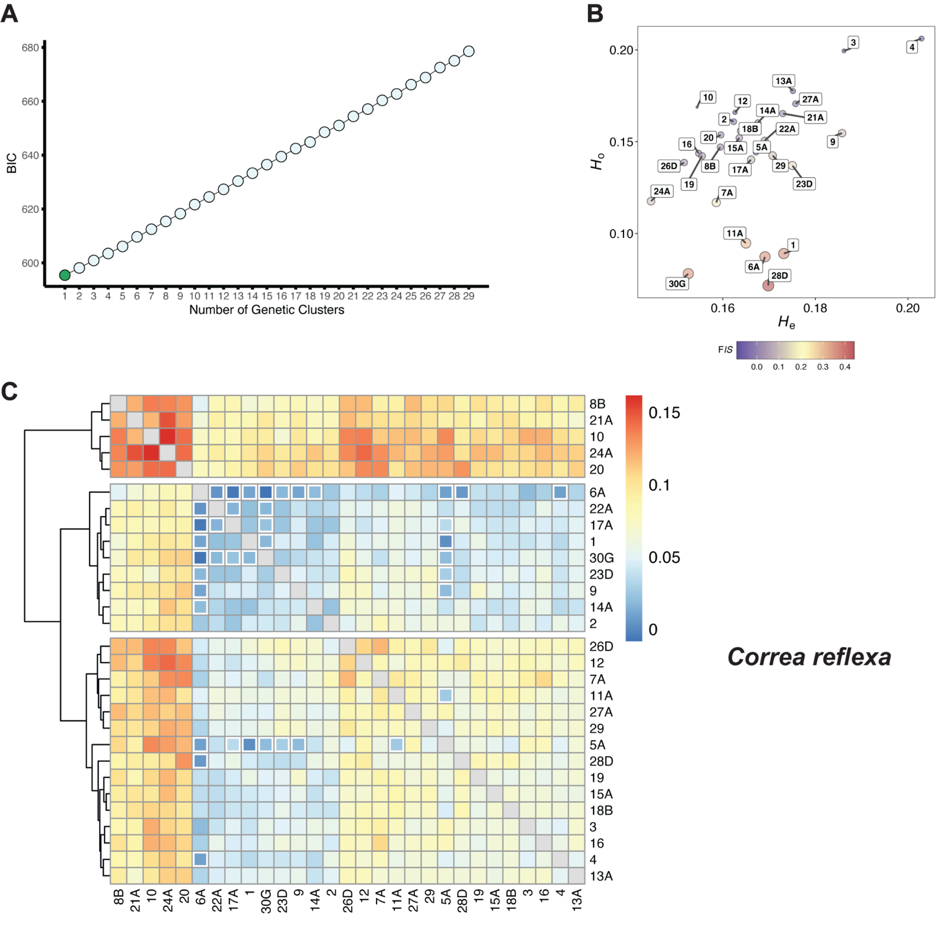

**Figure S4** (A) Modelling of the genetic differences among “populations” using a DAPC. (B) Comparison of genetic diversity parameters, where observed heterozygosity (*H*_O_; y-axis) was plotted as a function of expected heterozygosity (*H*_E_; x-axis). Locations are coloured by the overall inbreeding coefficient (*F*_IS_), and circle size is proportional to this value. (C) Heatmap of pairwise *F*_ST_ between all sampling location populations (Weir & Cockerham *F*_ST_). White frame squares reflect confidence intervals (CI) less than 0. In figures b) and c), the colour blue indicates lower estimates, and yellow and red indicate middle and higher estimates, respectively.

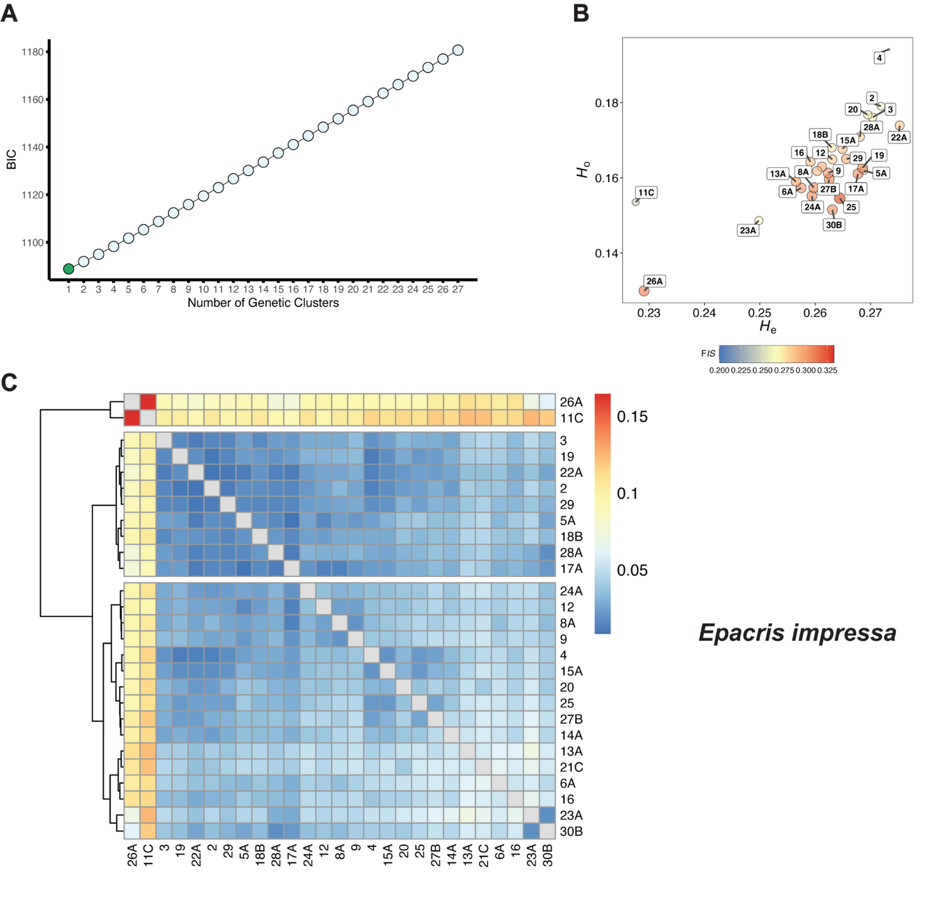

**Figure S5** (A) Modelling of the genetic differences among “populations” using a DAPC. (B) Comparison of genetic diversity parameters, where observed heterozygosity (*H*_O_; y-axis) was plotted as a function of expected heterozygosity (*H*_E_; x-axis). Locations are coloured by the overall inbreeding coefficient (*F*_IS_), and circle size is proportional to this value. (C) Heatmap of pairwise *F*_ST_ between all sampling location populations (Weir & Cockerham *F*_ST_). White frame squares reflect confidence intervals (CI) less than 0. In figures b) and c), the colour blue indicates lower estimates, and yellow and red indicate middle and higher estimates, respectively.

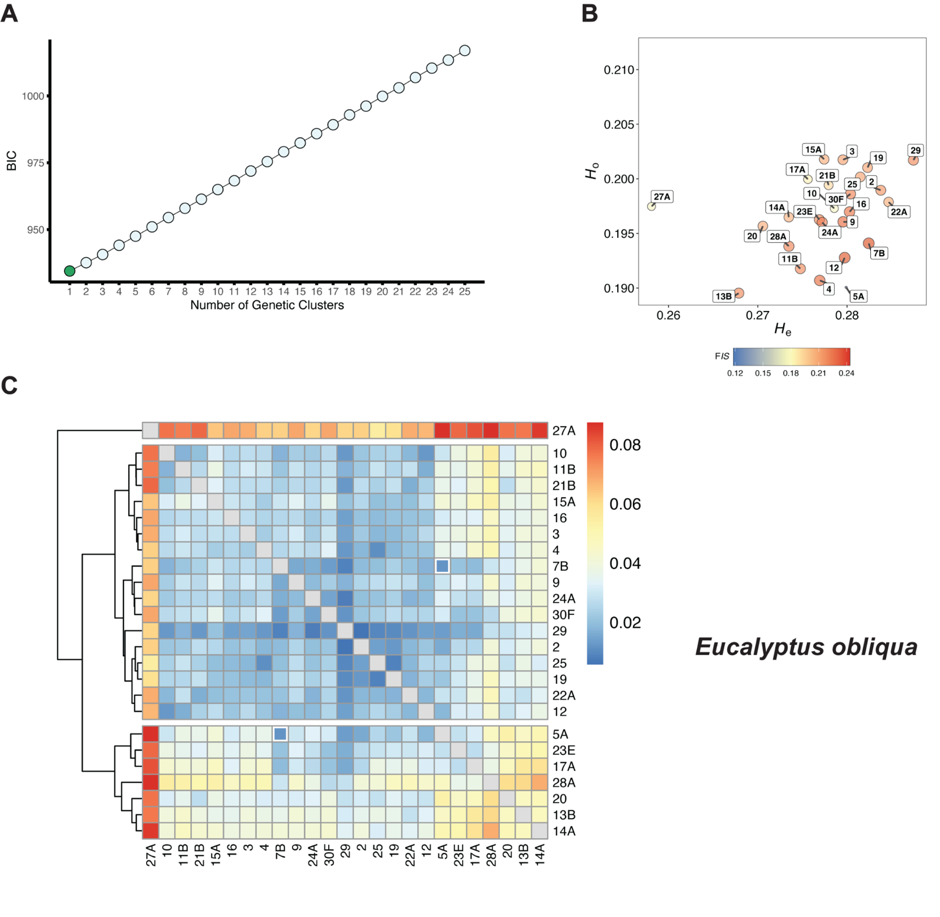

**Figure S6** (A) Modelling of the genetic differences among “populations” using a DAPC. (B) Comparison of genetic diversity parameters, where observed heterozygosity (*H*_O_; y-axis) was plotted as a function of expected heterozygosity (*H*_E_; x-axis). Locations are coloured by the overall inbreeding coefficient (*F*_IS_), and circle size is proportional to this value. (C) Heatmap of pairwise *F*_ST_ between all sampling location populations (Weir & Cockerham *F*_ST_). White frame squares reflect confidence intervals (CI) less than 0. In figures b) and c), the colour blue indicates lower estimates, and yellow and red indicate middle and higher estimates, respectively.

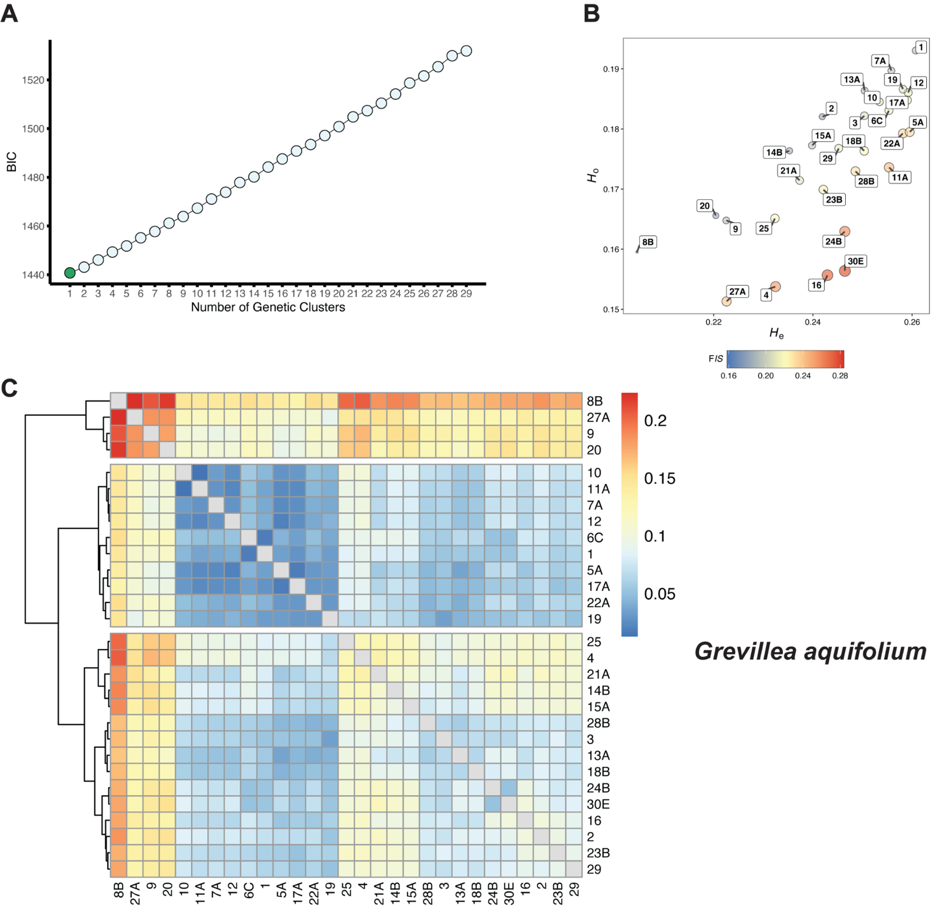

**Figure S7** (A) Modelling of the genetic differences among “populations” using a DAPC. (B) Comparison of genetic diversity parameters, where observed heterozygosity (*H*_O_; y-axis) was plotted as a function of expected heterozygosity (*H*_E_; x-axis). Locations are coloured by the overall inbreeding coefficient (*F*_IS_), and circle size is proportional to this value. (C) Heatmap of pairwise *F*_ST_ between all sampling location populations (Weir & Cockerham *F*_ST_). White frame squares reflect confidence intervals (CI) less than 0. In figures b) and c), the colour blue indicates lower estimates, and yellow and red indicate middle and higher estimates, respectively.

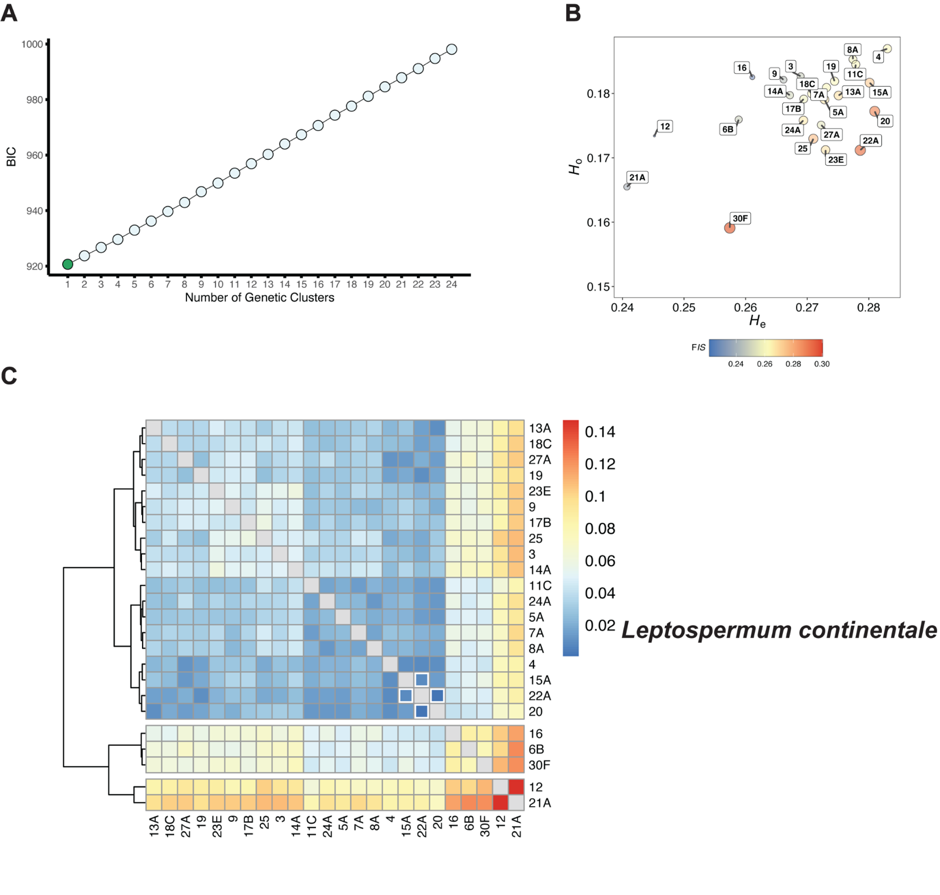

**Figure S8** (A) Modelling of the genetic differences among “populations” using a DAPC. (B) Comparison of genetic diversity parameters, where observed heterozygosity (*H*_O_; y-axis) was plotted as a function of expected heterozygosity (*H*_E_; x-axis). Locations are coloured by the overall inbreeding coefficient (*F*_IS_), and circle size is proportional to this value. (C) Heatmap of pairwise *F*_ST_ between all sampling location populations (Weir & Cockerham *F*_ST_). White frame squares reflect confidence intervals (CI) less than 0. In figures b) and c), the colour blue indicates lower estimates, and yellow and red indicate middle and higher estimates, respectively.

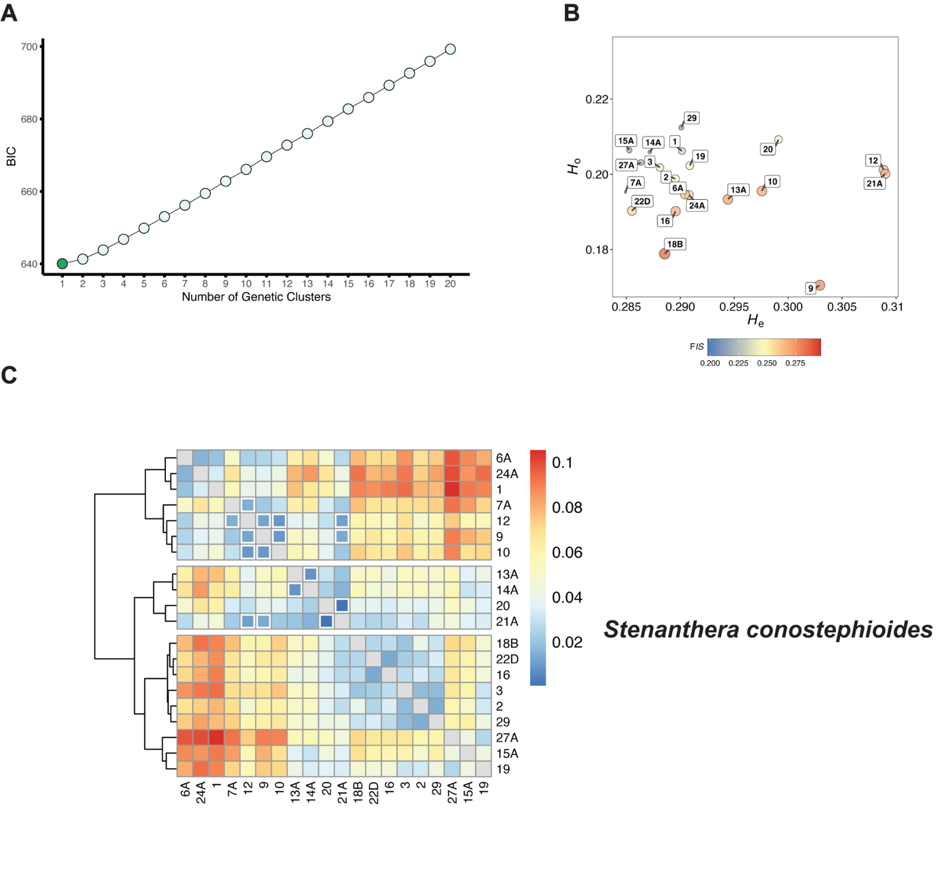

**Figure S9** (A) Modelling of the genetic differences among “populations” using a DAPC. (B) Comparison of genetic diversity parameters, where observed heterozygosity (*H*_O_; y-axis) was plotted as a function of expected heterozygosity (*H*_E_; x-axis). Locations are coloured by the overall inbreeding coefficient (*F*_IS_), and circle size is proportional to this value. (C) Heatmap of pairwise *F*_ST_ between all sampling location populations (Weir & Cockerham *F*_ST_). White frame squares reflect confidence intervals (CI) less than 0. In figures b) and c), the colour blue indicates lower estimates, and yellow and red indicate middle and higher estimates, respectively.

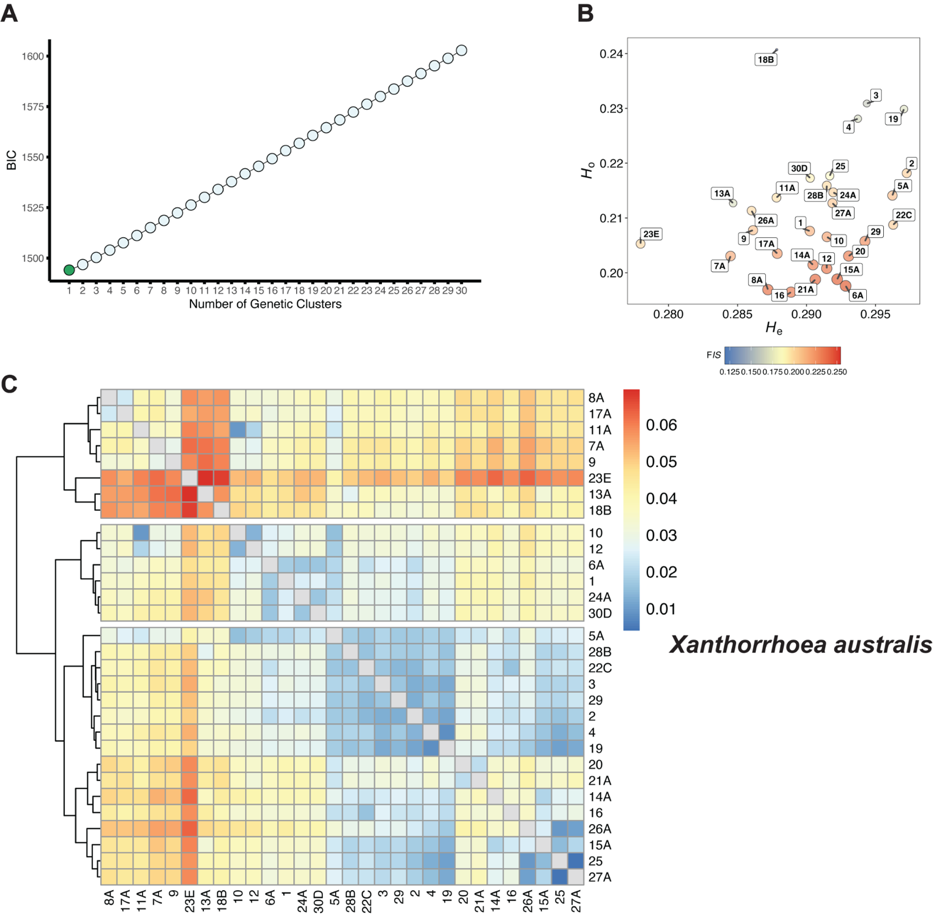
**Figure S10** (A) Modelling of the genetic differences among “populations” using a DAPC. (B) Comparison of genetic diversity parameters, where observed heterozygosity (*H*_O_; y-axis) was plotted as a function of expected heterozygosity (*H*_E_; x-axis). Locations are coloured by the overall inbreeding coefficient (*F*_IS_), and circle size is proportional to this value. (C) Heatmap of pairwise *F*_ST_ between all sampling location populations (Weir & Cockerham *F*_ST_). White frame squares reflect confidence intervals (CI) less than 0. In figures b) and c), the colour blue indicates lower estimates, and yellow and red indicate middle and higher estimates, respectively.

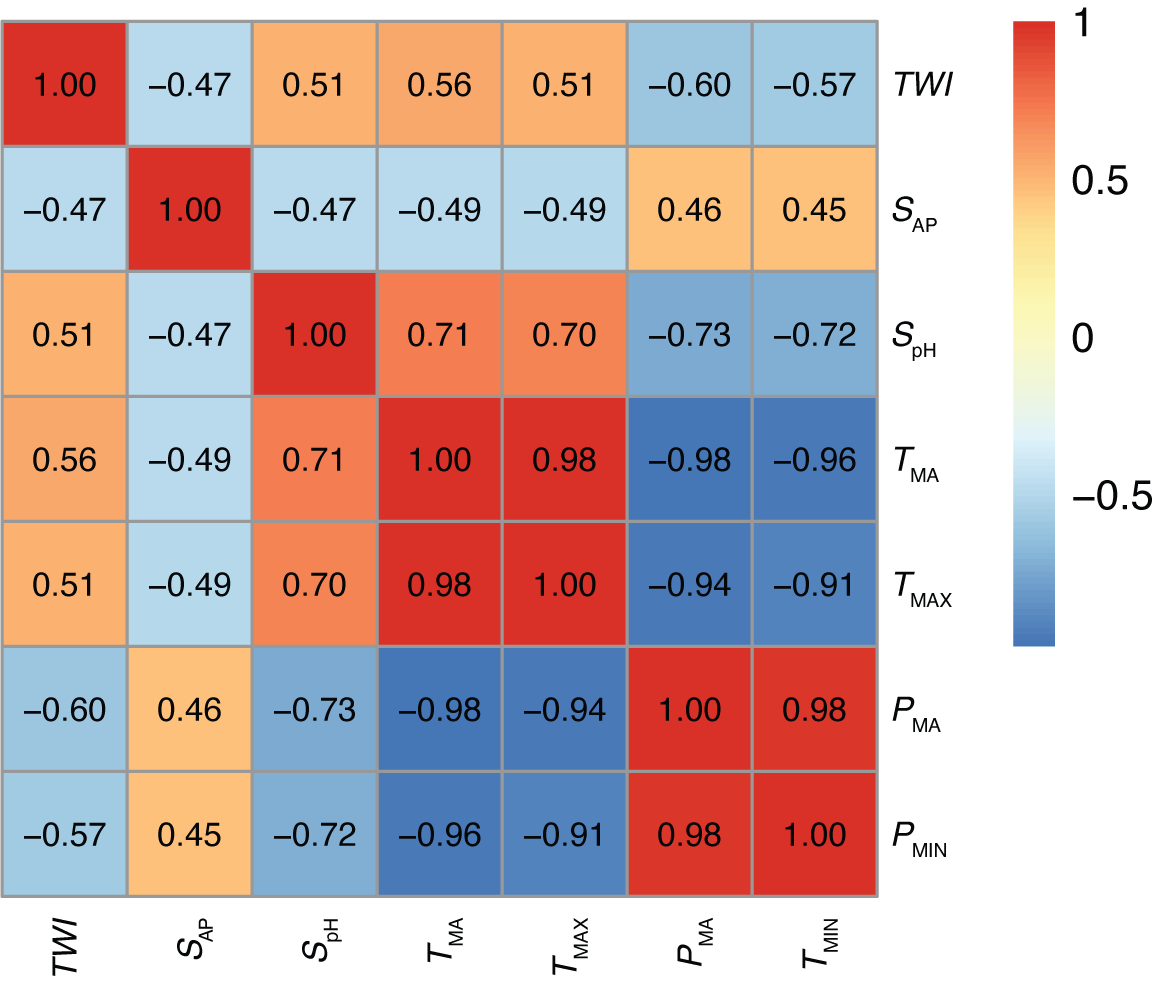

**Figure S11** Heatmap of Pearson correlation coefficients showing relationships between extracted environmental predictors across sampling locations for generalised dissimilarity modelling (GDM). Pearson’s values are displayed within each cell, ranging from strongly positive (red) to strongly negative (blue). Abbreviations of the variables are *TWI*–topographic wetness index, *S*_AP_–available phosphorus, *S*_pH_–soil pH in water and CaCl_2_, *T*_MA_–annual mean temperature, *T*_MAX_–maximum temperature of the warmest month, *P*_MA_–annual precipitation and *P*_MIN_–precipitation of driest month.

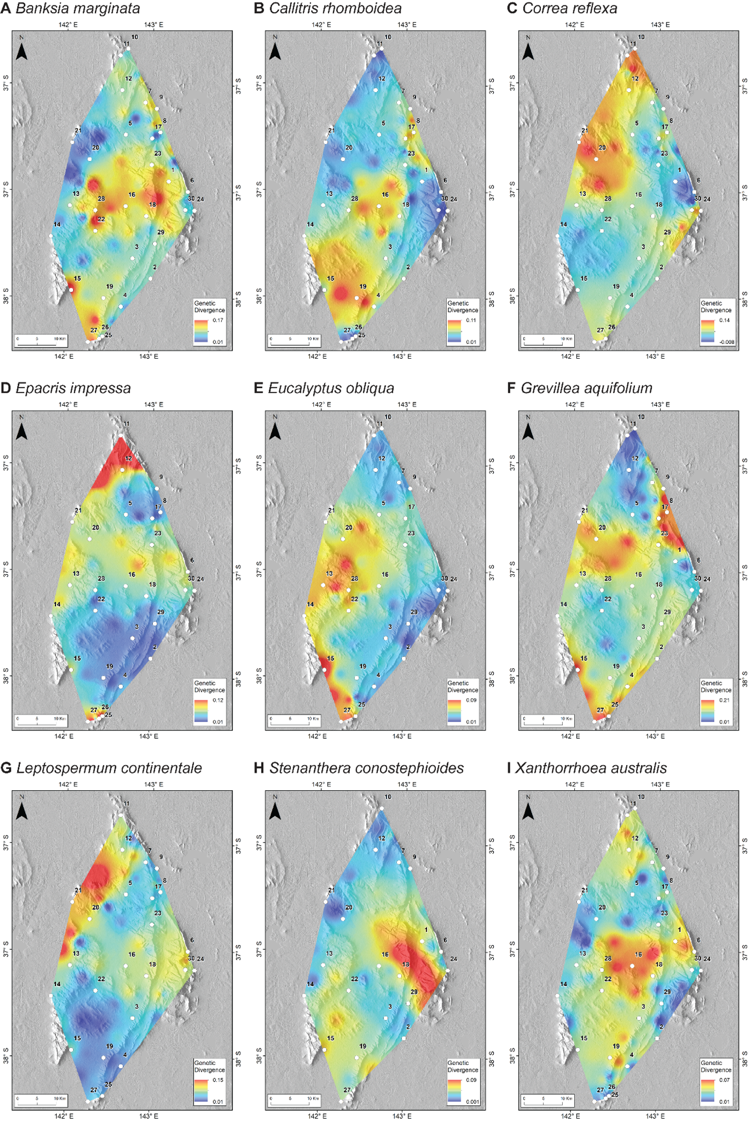

**Figure S12** Genetic divergence landscape of the nine species from Gariwerd (A–I). Colours indicate areas with low (blue) to high (red) species genetic divergence based on pairwise *F*_ST_ values. All other colours indicate the intermediate level of divergence. Small white circles indicate sampling locations (see Table S2 for location names).

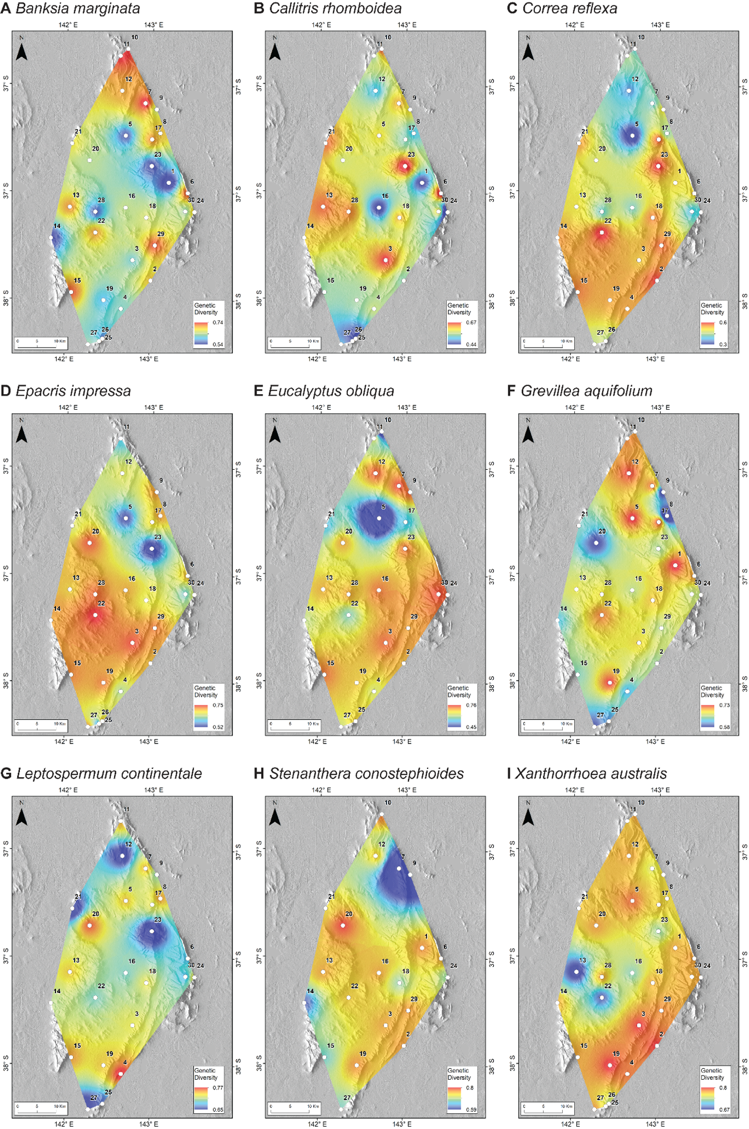

**Figure S13** Genetic diversity landscape of the nine species from Gariwerd (A–I). Colours indicate areas with low (blue) to high (red) species genetic diversity based on the proportion of polymorphic loci values. All other colours indicate the intermediate level of divergence. Small white circles indicate sampling locations (see Table S2 for location names).
